# Structures in multiple conformations reveal distinct transition metal and proton pathways in an Nramp transporter

**DOI:** 10.1101/403014

**Authors:** Aaron T. Bozzi, Christina M. Zimanyi, John M. Nicoludis, Brandon K. Lee, Casey H. Zhang, Rachelle Gaudet

**Author notes:** Co-first author (Aaron T. Bozzi, Christina M. Zimanyi).

## Abstract

Nramp family transporters—expressed in organisms from bacteria to humans—enable uptake of essential divalent transition metals via an alternating-access mechanism that includes proton co-transport. We present high-resolution structures of *Deinococcus radiodurans* (Dra)Nramp at complementary stages of its transport cycle to provide a thorough description of the Nramp transport cycle by identifying the key intramolecular rearrangements and changes to the metal coordination sphere. Strikingly, while metal transport requires cycling from outward-to inward-open states, efficient proton transport still occurs in outward-locked (but not inward-locked) DraNramp. We propose a model in which metal and proton enter the transporter via the same external pathway to the binding site, but follow separate routes to the cytoplasm, thus resolving the electrostatic dilemma of using a cation co-substrate to drive a cation primary substrate. Our results illustrate the flexibility of the LeuT fold to support a broad range of co-substrate coupling and conformational change mechanisms.

## Introduction

The <u>a</u>mino acid-<u>p</u>olyamine-organo<u>c</u>ation (APC) superfamily of secondary transporters encompasses a broad range of evolutionarily-related proteins that transport diverse substrates including neurotransmitters, metabolites, and transition metals in organisms throughout the tree of life (Vastermark et al., 2014; Wong et al., 2012). In humans alone, the APC superfamily encompasses 11 subfamilies of distinct solute carrier proteins (Perland and Fredriksson, 2017). These transporters harness the energy stored in preexisting transmembrane ion gradients. The LeuT fold (Yamashita et al., 2005) is the core structural unit that undergoes conformational rearrangements necessary for alternating access-based transport in the APC superfamily. This fold consists of ten transmembrane (TM) segments, divided into two pseudosymmetric, interlocking five-TM repeats, although many members have additional TMs. Primary substrates bind in a pocket formed by non-helical regions of TM1 and TM6, close to the center of the membrane. Co-transported coupling ions—typically Na^+^ and/or H^+^—bind at the interface between two proposed domains (Perez and Ziegler, 2013; Rudnick, 2013): a “bundle” formed by TMs 1, 2, 6, and 7; and a “scaffold” or “hash” domain comprising most or all of the remaining six TMs (Forrest and Rudnick, 2009). When all substrates are bound, conformational rearrangement closes an external vestibule between “bundle” and “scaffold” and opens an intracellular vestibule between the two domains to allow substrate release (Boudker and Verdon, 2010; Forrest et al., 2011; Shi, 2013). Despite the common fold, many APC members have little-to-no sequence identity, consistent with mechanistic divergences, including variance in the identity and stoichiometry of the coupled ions (Ma et al., 2012; Shaffer et al., 2009) and in which helices move the most to open and close the inner and outer gates (Kazmier et al., 2017; Kazmier et al., 2014a; Kazmier et al., 2014b; Krishnamurthy and Gouaux, 2012; Malinauskaite et al., 2014; Ressl et al., 2009; Shimamura et al., 2010; Weyand et al., 2008).

<u>N</u>atural <u>r</u>esistance-<u>a</u>ssociated <u>m</u>acrophage <u>p</u>roteins (Nramps) are APC-superfamily transition metal transporters that enable uptake of rare micronutrients such as Mn^2+^ in plants and bacteria and Fe^2+^ in animals (Cellier, 2012; Courville et al., 2006; Nevo and Nelson, 2006). Nramps bind and/or transport biologically essential divalent metals such as Mn^2+^, Fe^2+^, Co^2+^, Ni^2+^, Cu^2+^, Zn^2+^— and toxic metals like Cd^2+^, Pb^2+^, and Hg^2+^—but not the abundant alkaline earth metals Mg^2+^ and Ca^2+^ (Bozzi et al., 2016a; Ehrnstorfer et al., 2014). Metal uptake by Nramps involves proton co-transport, and many homologs also display considerable proton uniport—proton transport in the absence of added metal that suggests loose coupling between the co-substrates (Chen et al., 1999; Gunshin et al., 1997; Mackenzie et al., 2006; Nelson et al., 2002; Xu et al., 2004). Nramps have 11 or 12 TMs, the first ten forming a LeuT fold, as seen in structures of three bacterial Nramp homologs (Bozzi et al., 2016b; Ehrnstorfer et al., 2014; Ehrnstorfer et al., 2017), including our model system *Deinococcus radiodurans* (Dra)Nramp (Bozzi et al., 2016b). Conserved aspartate, asparagine, and methionine residues in TM1 and TM6 coordinate transition metal substrates as observed in an inward-open state (Ehrnstorfer et al., 2014), while only a metal-free outward-open state has been reported (Ehrnstorfer et al., 2017).

Here we provide the first complementary structures of the same Nramp homolog in multiple conformations, including the first metal-bound outward-open Nramp structure, and a novel inward-occluded structure. These allow us to fully illustrate the transport cycle for DraNramp. We also show that metal transport requires the expected alternating access bulk conformational change, whereas proton transport can occur via a more channel-like mechanism in the outward-open state. Using the structures and accompanying biochemical data, we delineate separate conserved transport pathways for metal and proton substrates and provide a mechanistic model encompassing substrate binding, release, and the conformational change process. We demonstrate novel modes of conformational rearrangement and ion shuttling in DraNramp compared to other LeuT-fold transporters, thus expanding the known repertoire of intramolecular dynamics and coupling mechanisms possible within this important protein family.

## Results

### Rational design and validation of conformationally-locked DraNramp constructs

A previously determined structure of a Fab-bound DraNramp in an inward-open conformation revealed the intracellular metal permeation pathway, or vestibule, between TMs 1a, 2, 5, 6b, 7, and 8 (Bozzi et al., 2016b). This structure was stabilized in an inward-open state by patches of mutations to intracellular loops 4-5, 6-7, and 10-11, and we thus refer to it as the Patch mutant. To observe additional conformational states of a transport cycle in a single Nramp homolog at high resolution, we developed two complementary conformationally-locked constructs for crystallization. Adding steric bulk along TM1a—for example a G45R mutation, which mimics a human anemia-causing mutation of a conserved glycine (Barrios et al., 2012)—prevented full closing of the intracellular vestibule and eliminated metal transport, emphasizing the importance of the alternating-access mechanism to DraNramp function(Bozzi et al., 2016b). Based on these findings we pursued the G45R mutant as a new inward-locked crystallization construct.

To develop a complementary outward-locked DraNramp construct, we adapted an approach previously described for the lactose transporter LacY (Kumar et al., 2014; Smirnova et al., 2013). By mapping extensive cysteine accessibility data onto the inward-open structure, we identified the external vestibule between TMs 1b, 6a, 3, 8, and 10 (Bozzi et al., 2016b). We created a panel of 11 tryptophan point mutants lining this predicted external vestibule (Figure 1A) to destabilize the inward-open state. An outward-locking mutation should severely impair metal transport, and indeed several mutants had impaired *in vivo* Co^2+^ uptake when expressed in *Escherichia coli* (Figure 1B and Figure 1—figure supplement 1A). We chose to pursue G223W—on TM6a one helical turn above the unwound metal-binding region—which like G45R eliminated Co^2+^ and Fe^2+^ metal transport (Figure 1—figure supplement 1B). Importantly, while a tryptophan modeled in the inward-open state at position 223 clashes with the top of TM10, the analogous glycine position in the recent structure of outward-open *Eremococcus coleocola* (Eco)Nramp (33% identity with DraNramp) lines a wide aqueous channel with adequate room for tryptophan’s bulk (Ehrnstorfer et al., 2017).

**Figure 1.**
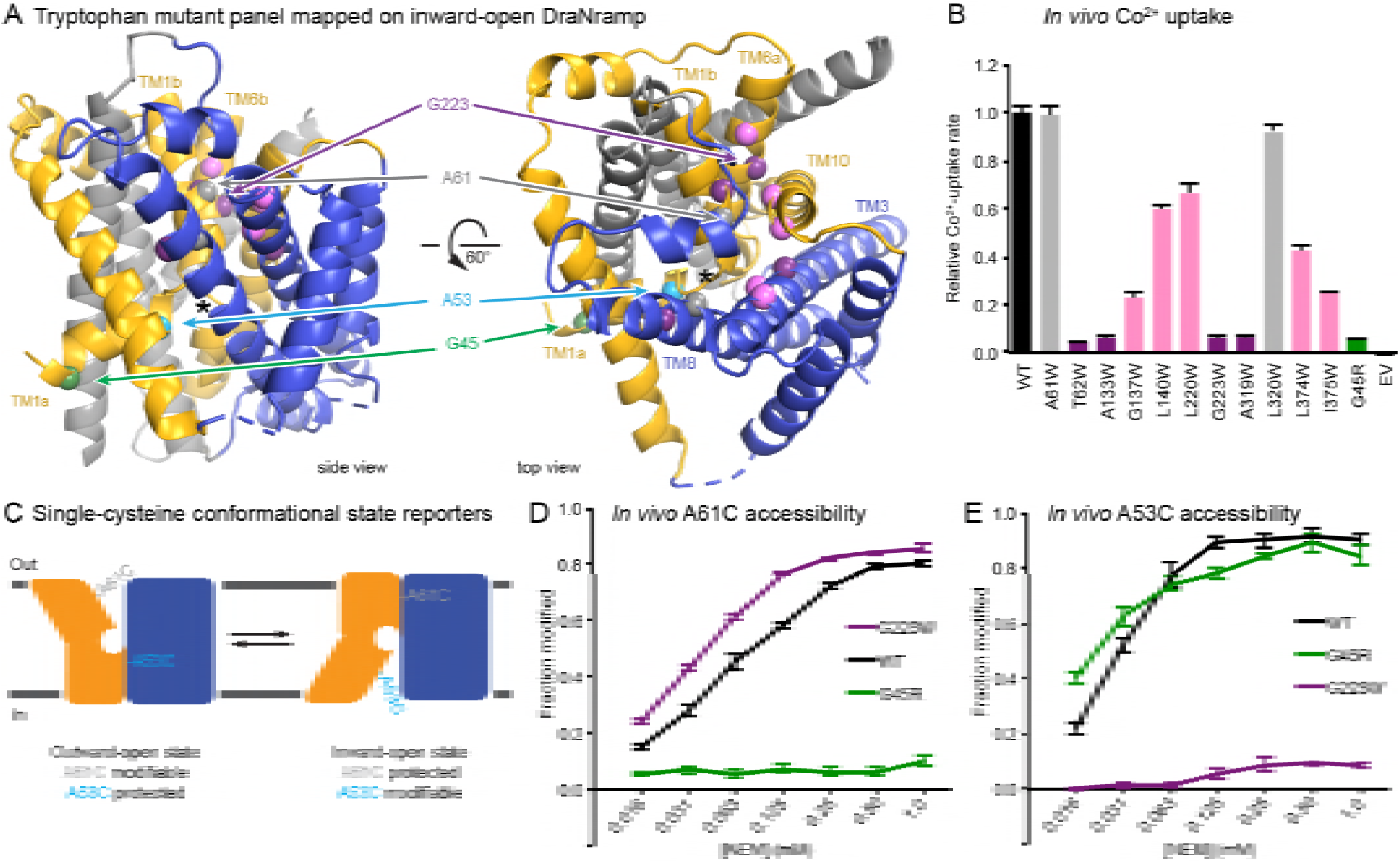
Design and validation of complementary conformationally-locked crystallization constructs. (A) Panel of 11 designed tryptophan mutants (pink, purple, and gray spheres; see (B)) and G45R disease-mutant mimic (green) mapped onto our initial inward-facing DraNramp structure and color-coded by their effect on transport activity in (B). Single-cysteine reporters A53C (cyan) and A61C (gray) are indicated. TMs 1, 5, 6, and 10 are colored gold, TMs 3, 4, 8, and 9 blue, and TMs 2, 7, and 11 gray. ^*^ marks the approximate location of the metal-binding site. (B) Relative *in vivo* Co^2+^ uptake rates for tryptophan mutant panel. Mutants colored gray, pink and purple did not affect, moderately decreased, or eliminated uptake, respectively. (C) Two complementary single-cysteine reporters enable assessment of the mutants’ conformational state preferences. For simplicity, in schematics DraNramp is depicted as two gold and blue domains that reorient to accomplish alternating access. (D) G45R prevented modification of the A61C outward-reporter, and (E) G223W fully protected the A53C inward-reporter, indicating that complementary conformational locking was achieved. Both reporters fully labeled the WT protein, likely because it cycles between both conformations during the assay. Data in B, D, and E are averages ± S.E.M. (n = 4). See also Figure 1—figure supplement 1.

To further validate this G223W construct, we measured bulk solvent accessibility of two single-cysteine reporters: A61C on TM1b, which is only exposed in DraNramp’s outward-open state (Bozzi et al., 2016b); and A53C on TM1a just below the metal-binding D56, a putative inward-open reporter based on comparing the Patch mutant and EcoNramp structures (Figure 1C and Figure 1—figure supplement 1C). WT-like DraNramp (with the indicated reporter cysteine and a C382S mutation to remove the lone endogenous cysteine) maintains a dynamic conformational equilibrium—even in the absence of added metal substrate—such that either reporter can be fully modified by the thiol-specific N-ethylmaleimide (NEM) at high concentrations (Figure 1D-E). G45R slightly increased A53C accessibility but fully protected A61C, indicating an inward-locked state, while G223W slightly increased A61C accessibility while fully protecting A53C, consistent with an outward-locked state (Figure 1D-E and Figure 1—figure supplement 1D). We have thus identified two complementary constructs that trap DraNramp in outward-locked (G223W) and inward-locked (G45R) states (Figure 2—figure supplement 2A). Using lipidic cubic phase (LCP) to mimic the hydrophobic membrane environment, we crystallized and determined the structures of G45R and G223W to resolutions of 3.0 and 2.4 Å, respectively, both significantly improved from our earlier DraNramp structure (3.94 Å) (Table 1 and Figure 2—figure supplements 1-2). The new high-resolution structures also allowed us to re-refine our original structure, including correction of a sequence registry error in TM11.

**Table 1.** DraNramp data collection and refinement statistics.

|  | **Inward-open apo**<br>**(Patch mutant + Fab)** | **Inward-occluded apo**<br>**(G45R)** | **Outward-open Mn<sup>2+</sup>-bound**<br>**(G223W ΔN34)** |
| --- | --- | --- | --- |
| **PDB ID** | 6D9W | 6C3I | 6BU5 |
| **SBGrid Data Bank ID** | 332, 333, 334 | 567 | 564 |
| **Data Collection** |  |  |  |
| Wavelength (Å) | 1.139 | 0.9793 | 0.9792 |
| Resolution range (Å) | 46.47 - 3.94 (4.08 - 3.94) | 50.00-3.00 (3.05-3.00) | 39.19-2.40 (2.49-2.40) |
| Space group | I222 | P2<sub>1</sub>2<sub>1</sub>2<sub>1</sub> | C121 |
| Unit cell (a, b, c) | 113.13, 132.08, 221.0 | 70.73, 97.85, 118.26 | 105.76, 80.39, 51.75 |
| Unit cell (α, β, γ) | 90.0, 90.0, 90.0 | 90.0, 90.0, 90.0 | 90.0, 94.72, 90.0 |
| Number of crystals | 3 | 2 | ~15 |
| Total reflections | 245867 | 66707 | 49321 |
| Unique reflections | 11791 (462) | 17206 (842) | 13983 (623) |
| Redundancy | 16.6 (11.4) | 3.9 (4.0) | 3.5 (1.7) |
| Completeness (%) | 95 (96) | 98.5 (98.8) | 82.1* (36.4) |
| Mean I/σ(I) | 6 (.61) | 6.41 (0.95) | 6.25 (1.56) |
| R<sub>merge</sub> | 0.171 | 0.223 (1.388) | 0.186 |
| R<sub>meas</sub> | 0.177 | 0.270 (1.608) | 0.210 |
| R<sub>pim</sub> | 0.047 | 0.133 (0.795) | 0.095 |
| CC<sub>1/2</sub> | 0.99 (0.189) | 0.993 (0.396) | 0.984 (0.584) |
| **Refinement** |  |  |  |
| Resolution range (Å) | 46.47 - 3.94 (4.12 - 3.94) | 30.0-3.0 (3.05-3.0) | 39.19-2.40 (2.486-2.40) |
| R<sub>work</sub> | 0.271 (0.253) | 0.249 (0.333) | 0.238 (0.276) |
| R<sub>free</sub> | 0.309 (0.343) | 0.270 (0.420) | 0.272 (0.374) |
| Number of atoms | 5764 | 5991 | 3213 |
| Protein | 5761 | 5812 | 3012 |
| Ligand | 3 | 150 | 139 |
| Water | -- | 29 | 62 |
| Protein residues | 786 | 774 | 398 |
| **Ramachandran plot** |  |  |  |
| Favored (%) | 754 (96.0) | 760 (98.2) | 389 (97.7) |
| Allowed (%) | 32 (4.0) | 14 (1.8) | 9 (2.3) |
| Outliers (%) | 0 (0.0) | 0 (0.0) | 0 (0.0) |
| RMS(bonds) | 0.004 | 0.004 | 0.004 |
| RMS(angles) | 1.02 | 0.97 | 0.92 |
| Average B-factor | 90.3 | 61.38 | 72.6 |
| Protein | 90.3 | 61.40 | 72.1 |
| Ligand | 197.0 | 66.01 | 86.4 |
| Water | -- | 32.82 | 63.8 |
| Number of TLS groups | 15 | 20 | 6 |

\*The G223WΔN34 dataset compiled from many small crystals is 99% complete to 3.4 Å, 95% complete to 3.0 Å, 90% complete to 2.85 Å, 80% complete to 2.7 Å, and 60% complete to 2.5 Å.

### Structure of G45R DraNramp reveals inward-occluded state

Unexpectedly, the G45R structure is not in an inward-open conformation as seen previously with our Fab-bound Patch mutant (Figure 2A) but instead adopts an inward-occluded, metal-free state (Figure 2B) that may represent an intermediate conformation between inward-open and outward-open states in the DraNramp transport cycle (Figure 2D). As in the inward-open apo state, the external vestibule remains sealed, with TM1b and TM6a forming tight hydrophobic packing with the tops of TM3 and TM10, and most TMs undergo little apparent displacement (Figure 2F-G). The major exception is TM1a, which swings ∼45° to partially seal the inward aqueous cavity in the G45R structure, a motion we previously showed to be essential to the transport cycle (Bozzi et al., 2016b). The intracellular ends of TM4 and TM5 also move slightly compared to their position in the inward-open state, further sealing the metal-binding site from the cytosol. Comparisons of the G45R and G223W structures indicate that, rather than preventing inward motion of TM1a as we had hypothesized (Bozzi et al., 2016b), the G45R mutation precludes TM4-TM5 from fully closing the inner gate, as any bulkier replacement for that absolutely-conserved glycine in our outward-open G223W structure would clash with E176 on TM5. Consequently, the intracellular vestibule to the metal-binding site is highly constricted yet there is no aqueous pathway to the binding site from the external side (Figure 2B). Structural alignments with the inward-open *Staphylococcus capitis* (Sca)Nramp (Ehrnstorfer et al., 2014) and outward-open EcoNramp (Ehrnstorfer et al., 2017) also indicate an intermediate conformation for the G45R structure, albeit closer to the inward-open state (Figure 2E and Figure 2—figure supplement 2D), confirming our assignment as inward-occluded.

**Figure 2.**
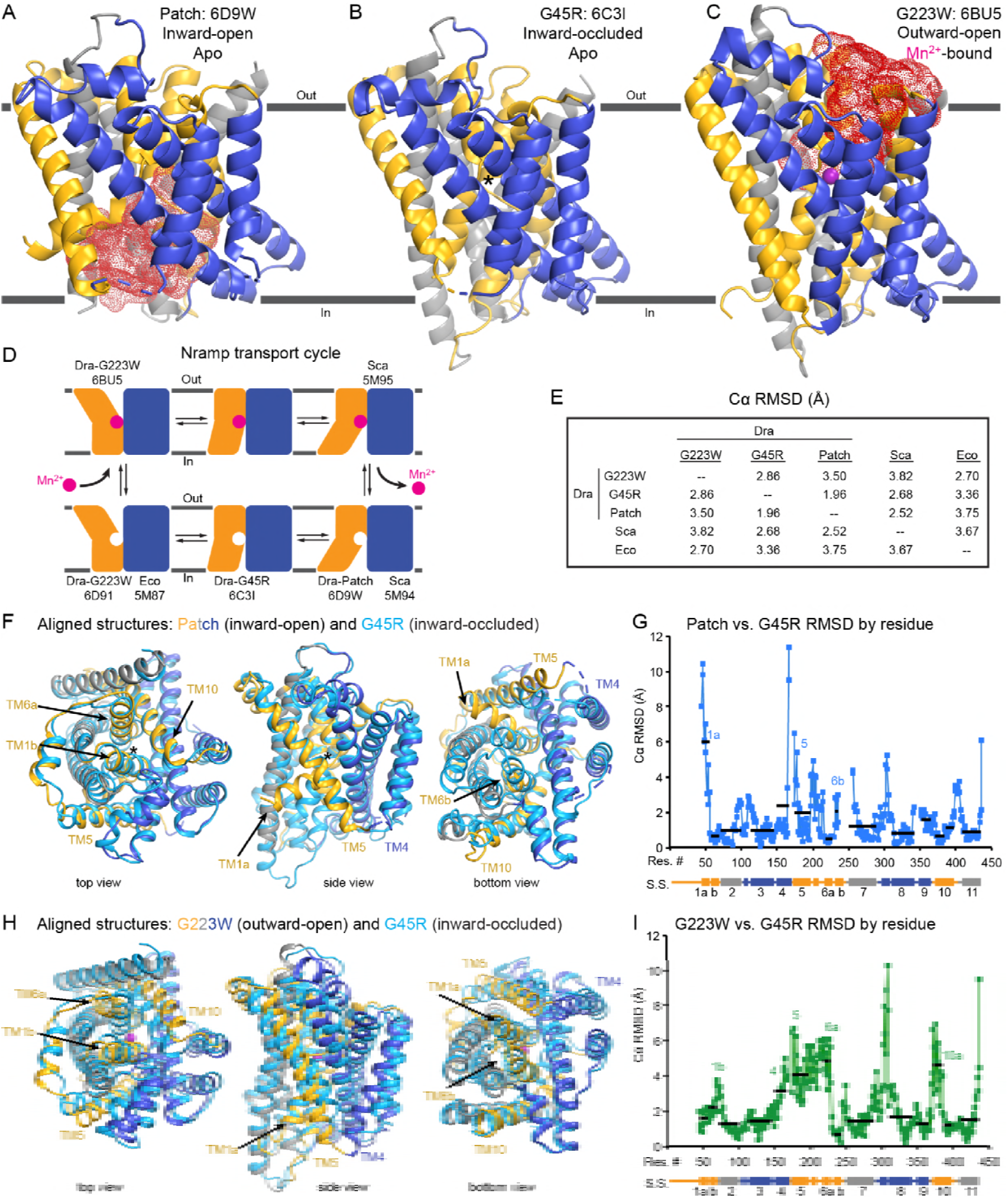
Crystal structures of DraNramp reveal two new conformations. (A) Updated DraNramp Patch mutant structure in an inward-facing apo state with a wide intracellular aqueous vestibule (red mesh) (Bozzi et al., 2016b). TMs 1, 5, 6, and 10 are colored gold, TMs 3, 4, 8, and 9 blue, and TMs 2, 7, and 11 gray. (B) The G45R structure revealed an apo inward-occluded conformation with no substantial extracellular or intracellular aqueous pathway to the metal-binding site, denoted by an ^*^ in structures where no metal is present. (C) The G223W structure revealed an outward-facing conformation with the physiological substrate Mn^2+^ (magenta sphere) bound at the bottom of a substantial extracellular aqueous cavity (red mesh). All structures are viewed from within the membrane. (D) The complete Nramp transport cycle likely consists of at least six distinct conformational states. Including the structures of *S. capitis* (Sca) (Ehrnstorfer et al., 2014) and *E. coleocola* (Eco) (Ehrnstorfer et al., 2017) homologs, we now have structures of five of these conformations. (E) Pairwise RMSD values for the superposition of the 355 Cα atoms present in all three DraNramp structures, and ScaNramp and EcoNramp. G223W superimposes best with the outward-facing EcoNramp, validating our mutagenesis strategy to obtain an outward-open conformation. G45R superimposes better with the inward-facing Patch mutant than outward-facing G223W, suggesting it represents an inward-occluded conformation. (F) Superposition of Patch mutant (gold, gray, and blue) and G45R (cyan) and (G) RMSD calculated at each Cα. These structures superimpose well except for TM1a. (H) Superposition of G223W (gold, gray, and blue) and G45R (cyan) and (I) RMSD calculated at each Cα. The most significant rearrangements involve TMs 1b, 4, 5, 6a, and 10. In panels G and I, black lines indicate average RMSD for each TM, with TMs 1, 6, and 10 divided into two halves. In panels F and H, the central view is rotated 45° along the vertical axis from the view in panels A-C, while the left and right views are 90° rotations of that central view. See also Figure 2—figure supplements 1 and 2.

### Structure of G223W DraNramp provides first metal-bound outward-open state

The G223W structure represents an outward-open, metal-bound state that superimposes best with the outward-open EcoNramp structure (Figure 2E and Figure 2—figure supplement 2E). As predicted, the exogenous tryptophan lines a periplasmic-facing aqueous cavity leading to a bound Mn^2+^ in the center of the transporter, with tight helix packing below precluding metal passage to the cytoplasm. We also determined a G223W apo structure (Table 1—table supplement 1), which lacks electron density attributable to metal substrate in the binding site (Figure 2—figure supplement 2C) but is otherwise similar to the metal-bound state (Cα RMSD = 1.08 Å); hence we used the metal-bound structure for all further analyses. Compared to the inward-open and inward-occluded structures, in the outward-open state TM1b, TM6a, and the top of TM10 are splayed open, and loop 1-2 is displaced by ∼4 Å, to form a wide aqueous pathway to the metal-binding site (Figure 2H-I). On the cytoplasmic side, TM4 and TM5 move significantly (by ∼8 Å) straddling TM8 and approaching TM1a, while TM1a also approaches TM8 to fully shut the interior aqueous vestibule (Figure 2H-I).

### Comparisons of DraNramp structures define a scaffold and flexible regions

Based on overall superpositions of the three DraNramp structures, TMs 1, 4, 5, 6, and 10 show the largest displacements to switch metal-binding site accessibility (Figure 2F-I). The remaining TMs (2, 3, 7, 8, 9, and 11) would thus form a “scaffold,” which adjusts to accommodate the more significant movements of the other five TMs (Video 1).

To more objectively compare the intramolecular rearrangements that occur during the transport cycle, we calculated difference distance matrices (Richards and Kundrot, 1988), averaged by TM, for each pair of structures (Figure 3). These matrices confirm that TMs 1, 4, 5, 6, and 10 undergo the most significant displacements relative to the rest of the protein between the different structures. But rather than moving as a rigid body such as proposed in the “rocking bundle” model for LeuT (Forrest and Rudnick, 2009), these five TMs are also significantly displaced relative to each other.

**Figure 3.**
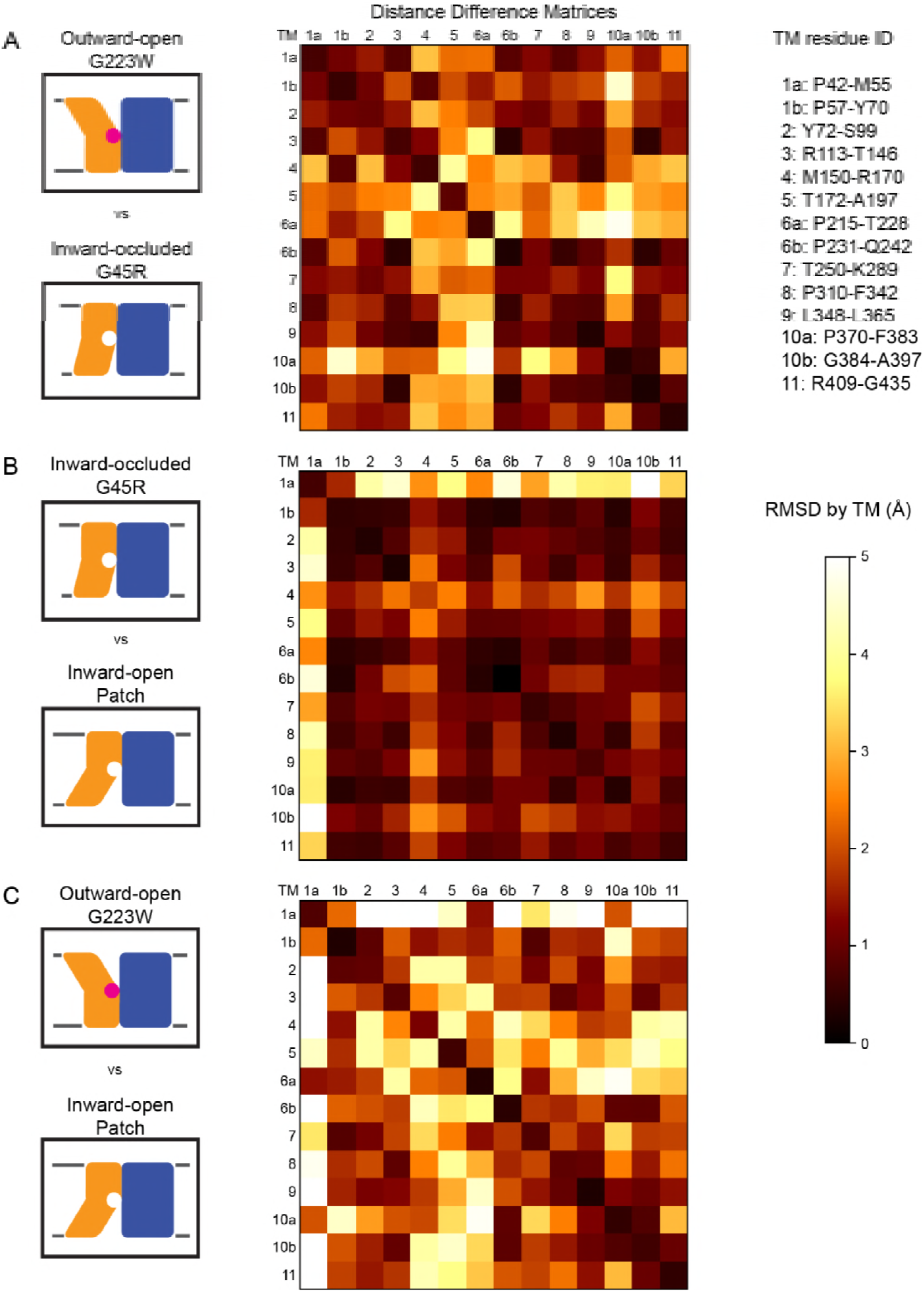
Distance difference matrices illustrate internal rearrangements during DraNramp conformational change. We calculated the pairwise Cα – Cα distances to obtain a 398 x 398 distance matrix for each of the three DraNramp structures. To compare two structures, we generated distance difference matrices(Richards and Kundrot, 1988), subtracting one distance matrix from the other. To simplify the data and focus on relative movements between helices, we then calculated the root-mean-squared deviations of submatrices, grouping residues within each helical segment. We used a total of 14 segments (top right), breaking TMs 1, 6, and 10 into two separate segments before and after their helix-breaking elements. The resulting 14 x 14 matrices compare (A) outward-open G223W and inward-occluded G45R, (B) inward-occluded G45R and inward-open Patch mutant, and (C) outward-open G223W and inward-open Patch mutant. Pairs of helices that remain stationary relative to each other have RMSD values close to 0 indicated by darker colors. In contrast, pairs of helices that rearrange significantly relative to each other have larger magnitude values indicated by lighter coloring in the heat map. These matrices show that (A) TMs 1b, 4, 5, 6a, and 10a undergo the greatest displacement relative both to the rest of the protein and to each other in the conformational change from inward-occluded to outward-open. (B) In contrast, the conformational change from inward-open to inward-occluded consists primarily of the large displacement of TM1a, with the rest of the protein remaining mainly stationary. (C) The comparison of the outward-open and inward-open states is essentially a sum of the two previous comparisons, with the large TM1a displacement added to the significant movements of TMs 1b, 4, 5, 6a, and 10a.

### The G223W outward-open structure reveals a metal-coordination sphere distinct from the inward-open state’s

Like other LeuT-family members, DraNramp relies on unwound regions of its TMs to bind substrates. A conserved DPGN sequence is non-helical in TM1—with the helix-breaking proline-glycine pair separating two metal-binding residues—while a conserved MPH sequence that includes the metal-binding methionine ends an unwound region in TM6 (Figure 4A-C). We used these two canonical motifs to generate an alignment of 6878 Nramp sequences (Figure 2—source data 1) and calculate the conservation of other important residues (Figure 2—figure supplement 1C). Interestingly, a third proline, P386 (83% conserved), enables the top of TM10 to swing to open the metal-binding site from the periplasm, while T228 (80%) on TM6 and N426 (99%) on TM11 stabilize the extended unwound TM6 region in the G223W structure (Figure 4—figure supplement 1A).

**Figure 4.**
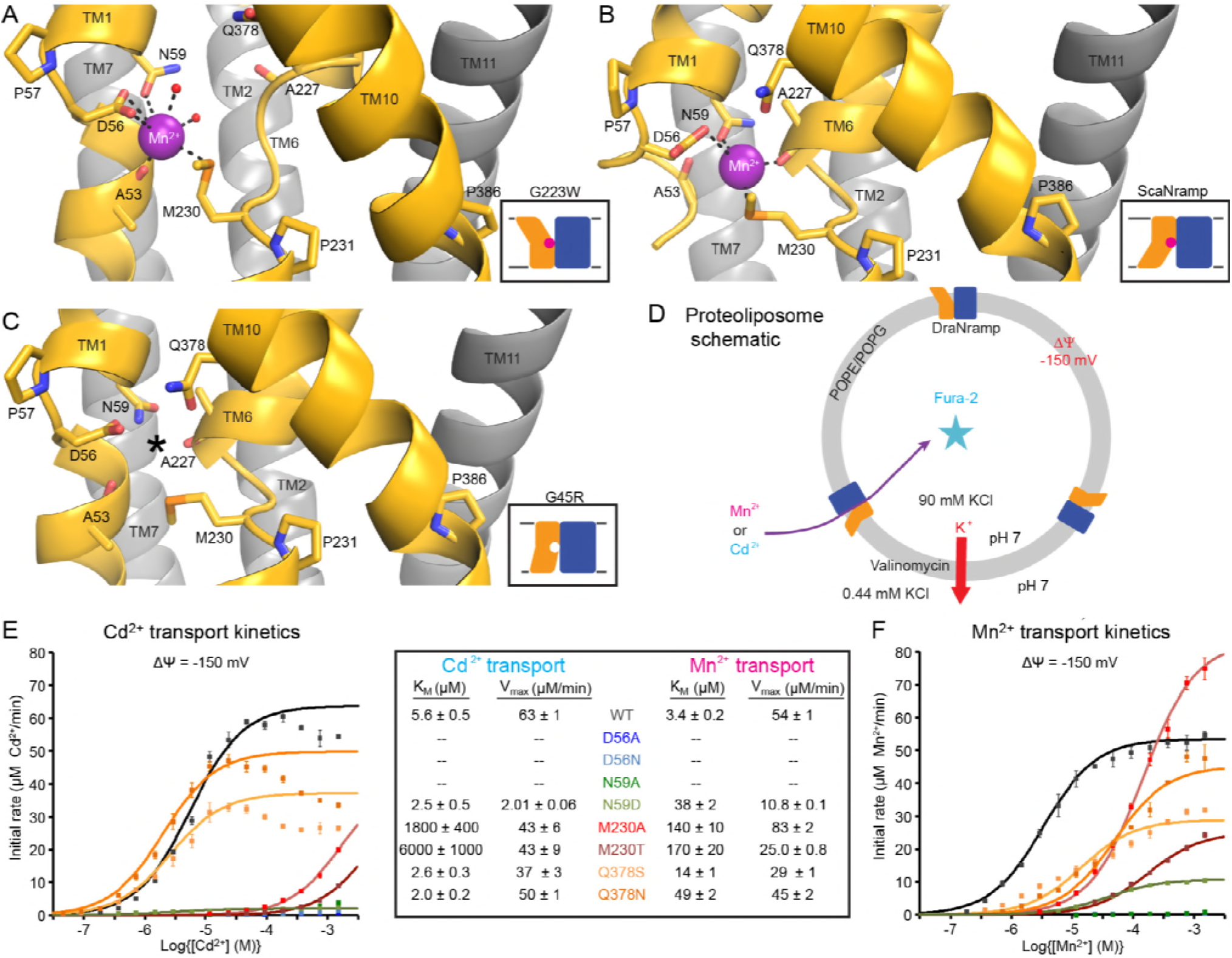
Metal-coordinating interactions vary across conformational states. (A) In the G223W outward-facing structure D56, N59, M230 and the A53 carbonyl, along with two water molecules, coordinate Mn^2+^. (B) In the inward-facing ScaNramp structure (PDB: 5M95, DraNramp residue numbering) D56, N59, M230, and the A227 carbonyl coordinate Mn^2+^. (C) In the G45R inward-occluded structure D56, N59, M230, Q378, and the A53 and A227 carbonyls are all close, suggesting that all six could simultaneously coordinate the metal substrate in a hypothetical similar Mn^2+^-bound conformation. Three conserved helix-breaking prolines on TM1 (P57), TM6 (P231), and TM10 (P386) confer the flexibility needed for the metal-binding site. The TM6 unwound region is extended in the outward-open state, while TM10 bends more dramatically at P386 in the inward-oriented states to close outside access to the metal-binding site. For clarity, TMs 3, 4, 5, 8, and 9 are omitted in panels A-C. (D) Schematic for liposome transport assay to measure Mn^2+^ or Cd^2+^ uptake. (E-F) Plots of initial rates vs. (E) Cd^2+^ or (F) Mn^2+^ concentration used to determine Michaelis-Menten constants (Table inset) for WT DraNramp and binding-site mutants. All mutants significantly impaired Mn^2+^ transport at lower [Mn^2+^], while only Q378 mutants did not severely impair Cd^2+^ transport. Data are averages ± S.E.M. (n = 3); errors on K_M_ and V_max_reflect uncertainty of the fit (shown as solid lines on the graphs) to the data. See also Figure 4—figure supplement 1.

The inward-open ScaNramp structure revealed a metal-binding site consisting of three conserved sidechains corresponding to D56, N59, and M230 in DraNramp, and a backbone carbonyl of A227 (Figure 4B) (Ehrnstorfer et al., 2014). In outward-open DraNramp, a Mn^2+^ binds both D56 and M230 (2.9 and 3.0 Å), with N59 slightly further away (3.4 Å) (Figure 4A). The increased unwinding of TM6a displaces the A227 carbonyl too far (6.5 Å) to coordinate the metal (Figure 4—figure supplement 1A-B). Instead, the A53 carbonyl coordinates the Mn^2+^ (2.0 Å)—our structure is thus the first to implicate this residue in the metal transport cycle. Two waters (2.7 and 2.8 Å) complete a Mn^2+^-coordination sphere. While the resolution remains too low to definitively define the coordination geometry, the electron density is consistent with Mn^2+^ interacting with both D56 oxygens and thus seven total ligands—rare but not unprecedented for Mn^2+^ (Barber-Zucker et al., 2017; Glasfeld et al., 2003). An ordered water network expands into the external vestibule as part of the extended metal coordination sphere (Figure 4—figure supplement 1A). A water is also tethered to the conserved H232 directly below the metal-binding M230, perhaps poised to hydrate the cation upon conformational change.

### The G45R inward-occluded structure suggests potential metal-binding role for conserved Q378

The inward-occluded G45R binding site contains no metal. The A53 carbonyl is farther from the other metal-binding residues than the A227 carbonyl (Figure 4C). This is consistent with a model in which Nramp metal transport involves a switch of ligands, perhaps with the A53 and A227 carbonyls both coordinating the metal substrate in an as-yet-uncaptured intermediate state.

In G45R, flexing of TM10 above its P386 pivot shifts Q378 (86%, 11% N) ∼5 Å to within hydrogen-bonding distance of metal-binding A227 and D56, perhaps stabilizing the negative charge during the empty transporter’s return to outward-open. While Q378 does not bind the metal substrate in either outward DraNramp (7.9 Å) or inward ScaNramp(Ehrnstorfer et al., 2014) (4.5 Å), its position in the G45R occluded intermediate suggests it may transiently bind during the transport process. Indeed, two independent molecular dynamics (MD) simulations of the inward-open ScaNramp showed a metal interaction with the Q378 oxygen (Bozzi et al., 2016a; Pujol-Gimenez et al., 2017), and mutations at this position impaired metal transport in HsNramp2 (Pujol-Gimenez et al., 2017).

To test the importance of the three conserved sidechains that coordinate Mn^2+^ (D56, N59, and M230; Figure 4A-B) and Q378 to metal transport, we purified a panel of mutants and reconstituted them into proteoliposomes (Figure 4D). D56A and D56N eliminated Mn^2+^ and Cd^2+^ transport (Figure 4E-F and Figure 4—figure supplement 1C-D), confirming the importance of D56. N59A severely and N59D moderately reduced transport of both metals (Figure 4E-F and Figure 4—figure supplement 1C-D). Both M230A and M230T transport both metal substrates (Figure 4—figure supplement 1C-D), but with lower apparent affinity than WT (Figure 4E-F). Consistent with our previous findings, removing M230—the lone sulfur-containing metal-binding residue—affects Cd^2+^ more than Mn^2+^ transport, reflecting the importance of the semi-covalent interaction Cd^2+^ can form with sulfur (Bozzi et al., 2016a). Lastly, Q378S and Q378N both preserved significant transport of both metals (Figure 4—figure supplement 1C-D). However, these mutants notably impaired transport at lower concentrations for Mn^2+^—but interestingly not Cd^2+^ (Figure 4E-F). That the native glutamine is essential for efficient transport of the biological substrate Mn^2+^ but dispensable for Cd^2+^ uptake (Figure 4—figure supplement 1E) suggests that the two metals interact differently with their surrounding ligands during the transport process, corroborating the differential effects of M230 mutations.

### DraNramp proton transport does not require large conformational change

We reconstituted G45R and G223W into proteoliposomes to assess their metal and proton transport (Figure 5A). Consistent with our *in vivo* findings, neither G45R nor G223W transport Mn^2+^, Cd^2+^, or Co^2+^ (Figure 5B and Figure 5—figure supplement 1A-B). Surprisingly, in the presence of a favorable negative membrane potential, G223W enabled larger basal H^+^ influx than WT, while G45R had no H^+^ flux (Figure 5C). Interestingly, while Mn^2+^ co-transport stimulated H^+^ uptake for WT, adding Mn^2+^ did not further stimulate either G45R or G223W (Figure 5C). These results suggest that Nramp metal and proton transport proceed via separate routes, with proton transport requiring only that the protein sample the outward-open state.

**Figure 5.**
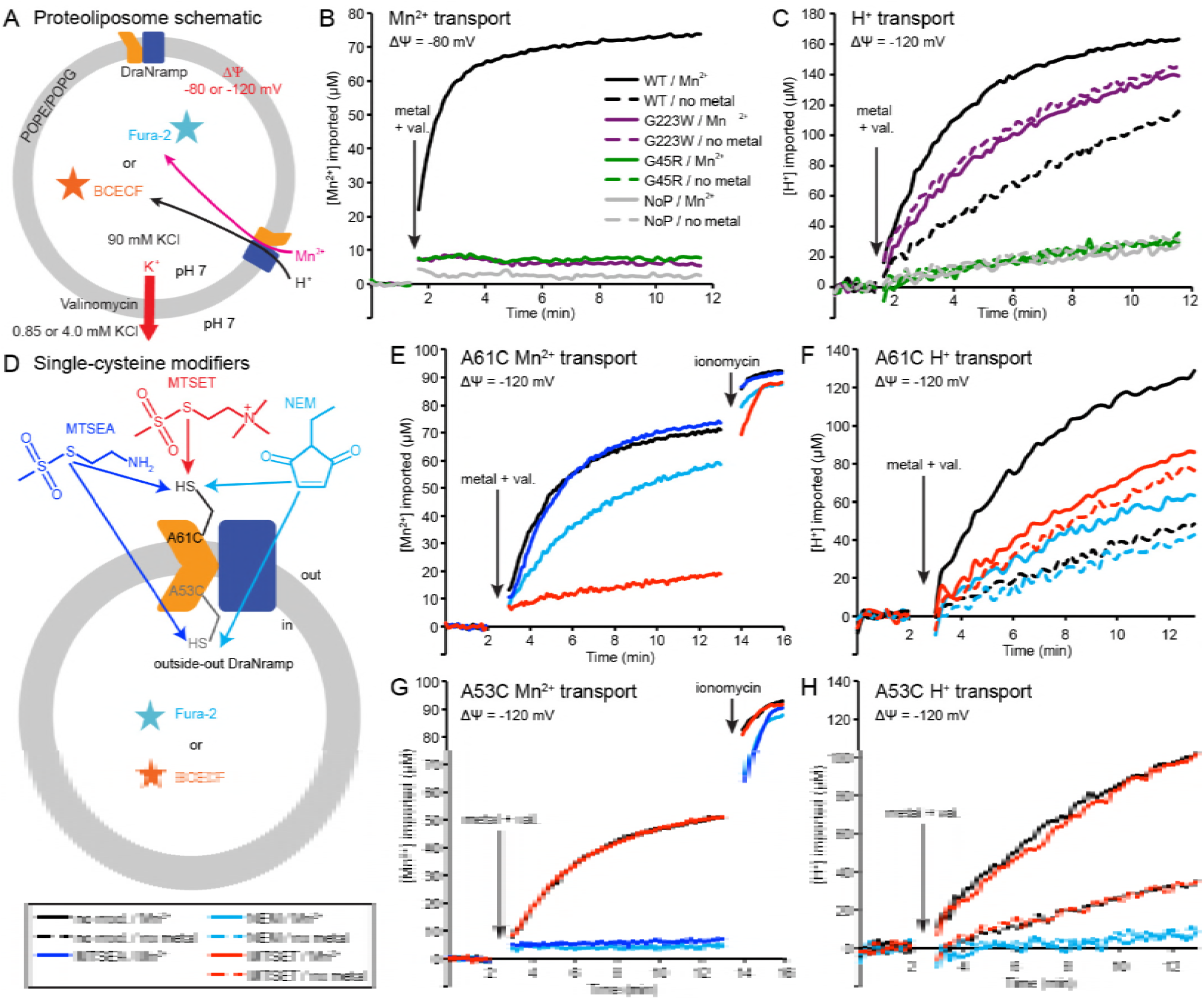
Outward-locked mutants retain H^+^ transport ability. (A) Proteoliposome assay schematic for monitoring M^2+^ and H^+^ import. Membrane potential (Δψ) was established using K^+^ gradients and valinomycin. (B) Mn^2+^ uptake over time; WT enabled robust uptake while G45R and G223W showed no activity. NoP = No Protein control liposomes. (C) H^+^ uptake over time; WT enabled significant H^+^ uptake without metal (H^+^ uniport), further stimulated by Mn^2+^. G223W showed a larger H^+^ uniport rate, but Mn^2+^ provided no enhancement; G45R showed no H^+^ transport activity. Traces are representative of four experiments. (D) Schematic for *in vitro* cysteine modification of A53C (inside accessible) and A61C (outside accessible). MTSEA and NEM are membrane permeable, whereas permanently-charged MTSET is not. (E,G) Mn^2+^ and (F,H) H^+^ transport over time in the presence or absence of cysteine modifying agents. Adding ionomycin allows divalent cation entry to achieve maximum signal. Traces are representative of three experiments. See also Figure 5—figure supplement 1.

To test this new hypothesis, we reconstituted the A53C and A61C mutants, with single cysteines located just below or above the metal-binding site respectively (Figure 5—figure supplement 1D). Both retain significant metal transport (Figure 5E,G and Figure 5—figure supplement 1E-F), and can be targeted with cysteine-specific modifiers to post-translationally add bulky and/or charged wedges to impede conformational change (Figure 5D and Figure 5—figure supplement 1C). Charged, and thus membrane-impermeable, MTSET nearly eliminated metal transport by A61C, while uncharged NEM or MTSEA moderately impaired or did not affect transport, respectively (Figure 5E and Figure 5—figure supplement 1E), a result consistent with our previous *in vivo* findings that adding steric bulk, but not formal charge, is tolerated at this position (Bozzi et al., 2016b). In addition, MTSET-treated A61C replicated G223W’s H^+^-transport behavior, with higher H^+^ uniport compared to unmodified transporter, but little stimulation by Mn^2+^ (Figure 5F). In contrast MTSET had no effect on Mn^2+^ and Cd^2+^ transport by A53C, but membrane-permeable MTSEA and NEM both eliminated transport (Figure 5G and Figure 5—figure supplement 1F). Regarding H^+^ uptake, MTSEA-and NEM-treated A53C resembled G45R, with no basal uniport or Mn^2+^ stimulation, while again unmodified and MTSET-treated A53C both behaved similarly to WT (Figure 5H). These findings show that essentially all activity in proteoliposomes, which likely contain a mix of inside-out and outside-out transporters, comes from outside-out DraNramp. MTSET, which should inhibit any inside-out A53C, did not affect transport, indicating that inside-out transporters contribute negligibly to the total activity, a result we more fully explore in an accompanying paper (Bozzi et al., 2018). Consistently, while MTSET treatment spares inside-out A61C from labeling, it nevertheless nearly eliminated metal transport.

In summary, these experiments with permanently-locked crystallization constructs or chemically-locked cysteine mutants demonstrated that while metal transport requires complete conformational cycling, proton transport does not require large-scale conformational change and can proceed through DraNramp’s outward-open state but not its inward-open state.

### Conserved salt-bridge network provides potential proton pathway to cytoplasm

Our *in vitro* results suggested that proton transport occurs via a pathway separate from the intracellular metal-release vestibule, which remains closed to bulk solvent in the proton-transporting G223W mutant. Below the metal-binding site begins a network of highly-conserved hydrophilic residues, including at least seven potentially protonatable sidechains, that leads from the metal-binding D56 through a tight corridor between TMs 3, 4, 8, and 9 to the cytoplasm (Figure 6A-C). In contrast to the external and intracellular vestibules proposed as metal entrance and release pathways, the helices and residues within this polar network undergo little rearrangement between the three DraNramp structures, except the intracellular end of TM4 (Figures 2F-I and 3). Highly-conserved residues surrounding the metal-binding D56 include H232 on TM6b (100% as defined) and E134 (98%, TM3)—which have each been proposed as the Nramp proton transfer point(Ehrnstorfer et al., 2017; Pujol-Gimenez et al., 2017). Across from E134 lies a conserved salt-bridge pair: D131 (TM3, 93%) and R353 (TM9, 78%). Approximately 9 Å below, a second conserved salt-bridge, E124-R352 (94% and 87%), links the same two helices. This network could provide the route for proton uniport in the outward-open conformation.

**Figure 6.**
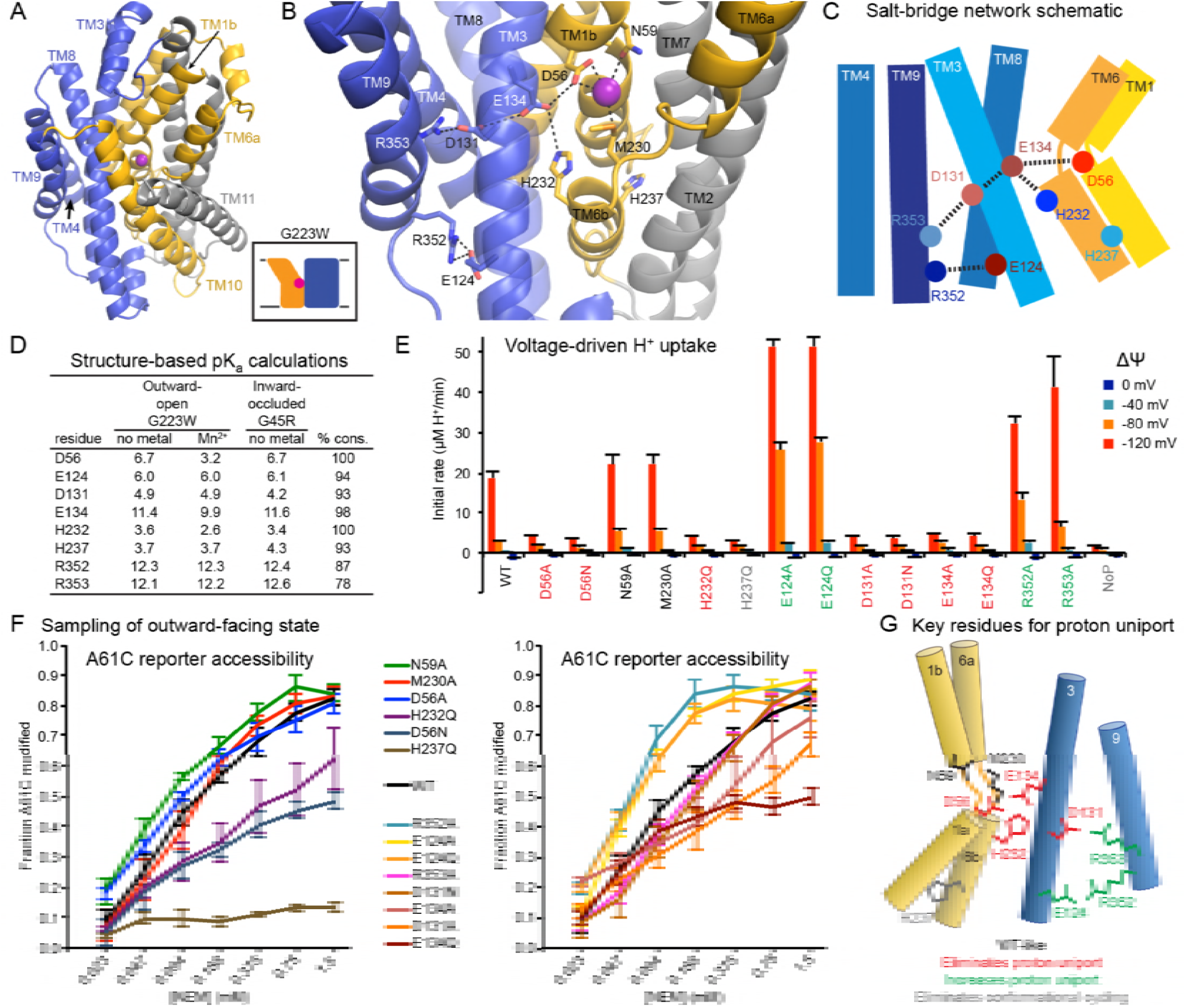
Conserved salt-bridge network provides pathway for proton uniport. (A) View from an angle above the membrane looking down into the extracellular aqueous cavity in the outward-facing G223W structure. (B) Zoomed-in view (with TM10 and TM11 omitted for clarity) and (C) schematic showing a network of conserved protonatable residues that originates from the metal-binding site and extends into the cleft between TMs 3, 4, 8, and 9. H232 and E134 abut metal-binding M230 and D56, providing a connection to the D131-R353 and R352-E124 salt-bridge pairs. (D) pK_a_ estimates by PROPKA (Dolinsky et al., 2004) from high-resolution DraNramp structures. (E) Initial rates of basal H^+^ uptake at various Δψs (averages ± S.E.M.; n ≥ 4). Mutations to D56, E134, H232, and D131 eliminated H^+^ uniport observed for WT. N59 and M230 mutants retained significant H^+^ uptake. Mutations to E124, R352, and R353 enhanced H^+^ uniport. (F) Dose-response curve of outward-facing reporter A61C modification versus NEM concentration. All mutants except H237Q sampled the outward-facing state, which is required for H^+^ transport to occur (Figure 5). Data are averages ± S.E.M. (n ≥ 3). (G) Salt-bridge network schematic shows clustering of four residues required for H^+^ uniport (red) as well as three residues that may restrain this flux (green). See also Figure 6—figure supplement 1.

### Two conserved aspartates anchor the DraNramp proton transport pathway

To assess whether these residues could be proton carriers, we calculated predicted pK_a_ values for our outward-open and inward-occluded structures (Figure 6D) (Dolinsky et al., 2004). Surprisingly, D56 is the only residue with a pK_a_ in the ideal 6-7 range to facilitate proton exchange at a typical external pH. About 4 Å from D56, E134’s high pK_a_ indicates a near permanently-protonated state, while H232—4 Å below E134 and 7 Å from D56—has too low a pK_a_ to easily protonate, as does H237 further down TM6b. While E134 and H232 have separately been suggested as the Nramp proton-binding site (Ehrnstorfer et al., 2017; Pujol-Gimenez et al., 2017), our pK_a_ predictions suggest otherwise, as maintaining a formal change, especially on a histidine, would not be favorable in the protein core, while the E-to-Q mutant at that position in EcoNramp maintained WT-like proton transport ability (Ehrnstorfer et al., 2017), which argues against that residue as an essential transfer point. Within the TM3-TM9 salt-bridge network, R352 and R353 are likely protonated and positively charged, while their respective partners E124 and D131 are likely deprotonated and negatively charged. The predicted pK_a_ values of D131 and E124 indicate their amenability to protonation. Indeed, as D56’s predicted pK_a_ drops to 3.2 with Mn^2+^ bound, D131 becomes the best candidate to receive a proton.

We observed three distinct voltage-driven H^+^ transport phenotypes within a panel of mutants to highly-conserved residues (Figure 6E and Figure 6—figure supplement 1). First, removing either metal-binding residue N59 or M230 has little effect. Second, neutralizing any member of the D56-E134-H232-D131 network or the H237Q mutation drastically reduced H^+^ transport. Third, mutating any of E124, R352, R353—farthest from D56—increased H^+^ uniport across multiple voltages.

Outward-reporter A61C accessibility (Figure 1C) is consistent with each mutant sampling the outward-open state needed for proton transport (Figure 5), ruling out a conformation-locking explanation for the loss-of-function mutants. While some mutations perturbed the transporter’s conformational preference, A61C remained at least somewhat accessible in all cases except H237Q (Figure 6F).

In summary, D56 is the likely initial protonation point, with E134 and H232 positioned to chaperone the proton transfer to D131, while R352, R353, and E124 restrain this process (Figure 6G).

## Discussion

We propose a structure-based model for conformation cycling in DraNramp (Figure 7A-B). Starting from the outward-open state seen in our G223W structure (Figure 7A, left panel), metal binding and resulting proton entry into its release pathway may trigger bulk conformational rearrangement (see below for details). To close the external vestibule, TM6a, TM10, and to a lesser extent TM1b move inward above their respective non-helical hinge regions, with the TM6a movement propagated through the TM5-6 linker, causing rigid rotation of TM5 to begin to open the inner gate. From this transient occluded conformation similar to our G45R structure (Figure 7A, middle panel), additional movement of TM4-TM5 allows TM1a to bend upward to fully open the inner gate, enabling solvent access to and release of the metal, as the protein achieves a state similar to the Patch mutant DraNramp structure (Figure 7A, right panel) (Bozzi et al., 2016b). Analogously, to return to the outward-open state and complete the transport cycle, TM1a swings in to reach a conformation similar to G45R, then TM4-TM5 fully close on TM1a to seal the cytoplasmic vestibule while TM1b, TM6a, and TM10 separate to open the external vestibule.

**Figure 7.**
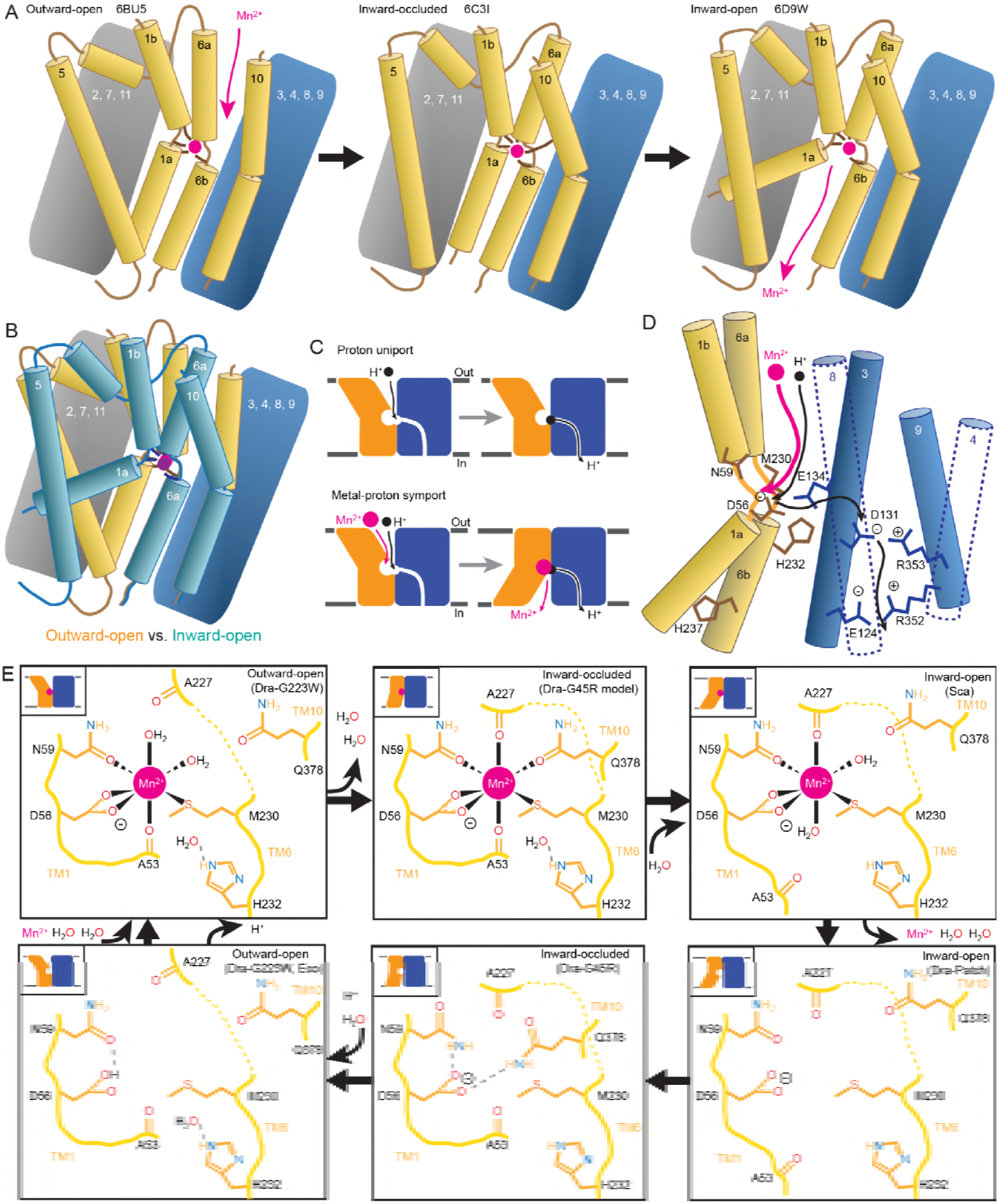
Mechanistic models for conformational change, proton transport, and metal transport. (A) Model of conformational change: The metal ion reaches the outward-facing binding site via the vestibule between TMs 1b, 6a, 10, 3, and 8. TM6a and 10 bend inward to close the vestibule in the inward-occluded conformation. TM6a’s inward bend pulls on TM5 via extracellular loop 5-6 to initiate opening of the inner gate. Finally, TM1a swings away from TM6b to open the intracellular vestibule into which the metal diffuses. (B) Superposition of outward-(gold) and inward-facing (teal) conformations illustrate key movements by TMs 1, 5, 6, and 10. (C) Overall transport model: While metal transport requires complete conformational cycling, proton uniport occurs through the outward-facing state. (D) Proton transport model: A proton transits the external vestibule to reach the binding site near the membrane center and initially binds to D56. Metal enters through the same passageway, ejecting the proton, which then passes to D131, with H232 and E134 facilitating the transfer. The proton ultimately reaches the cytoplasm through the polar network between TMs 3, 4, 8, and 9, while the metal must await a bulk conformational change that opens a separate pathway between TMs 1a, 2, 5, 6b, 7, and 8. Proton uniport follows the same route, with the D56-D131 transfer occurring at a slower rate without metal substrate. (E) Model for metal coordination during the transport process: The metal initially binds in the outward-facing state to D56, N59, M230 sidechains and the A53 carbonyl, shedding all but two water ligands. As the outward metal-permeation pathway closes, Q378 and the A227 carbonyl displace the waters so that the metal is fully coordinated by six amino acids in the occluded state. As the inner gate opens, Q378 and the A53 carbonyl withdraw and are replaced by waters. The metal is then released into the cytoplasmic aqueous vestibule. To facilitate the return to the outward-facing state, Q378 and N59 donate hydrogen bonds to negatively-charged D56 in the occluded return conformation seen in our G45R structure. Finally, as the transporter returns to an outward-open state, it may bind a proton in preparation for metal-binding.

Our *in vitro* assays showed that while DraNramp metal transport requires sampling of both outward-and inward-open states, proton uniport occurs in sterically outward-locked constructs. This supports a model where protons and metal travel through distinct pathways on the cytoplasmic side of the protein (Figure 7C-D), such that proton uniport is a feature of DraNramp’s outward-open state, whereas metal transport requires bulk rearrangement. In contrast, both protons and metal likely enter through the same aqueous pathway, as inward-locked proteins do not transport either substrate. Proton uniport occurs via a network of conserved protonatable residues leading from D56 in the metal-binding site to D131 in a salt-bridge network between TMs 3, 4, 8, and 9. This proton pathway is accessible in the outward-open state, thus enabling the well-documented proton uniport (Chen et al., 1999; Gunshin et al., 1997; Mackenzie et al., 2006; Nelson et al., 2002; Xu et al., 2004). The predicted protonation of D56 and subsequent transfer to D131, mediated through an E134/H232-stabilized transition state, may thus serve to restrain H^+^ entry. For metal-proton co-transport, Mn^2+^ interaction with D56 presumably stimulates proton transfer into the same salt-bridge network. We further explore the mechanistic details of DraNramp “proton-metal coupling” in an accompanying paper (Bozzi et al., 2018).

In our G223W structure, two water molecules coordinate Mn^2+^: one lies between the metal and A227’s carbonyl, the other the metal and Q378 (Figures 4A and 7E). We propose that after Mn^2+^ binds to D56, M230, A53, and N59 as in our G223W structure, the A227 carbonyl and Q378 both displace the two waters as the outer gate closes. DraNramp would thus reach a fully dehydrated metal-bound state not yet visualized but which may resemble our apo G45R inward-occluded structure (Figure 7E). Next, as the inner gate opens, the A53 carbonyl would exchange with a nearby water—such as the one bound to H232 in our G223W structure—as would Q378, to yield an inward-open metal-bound state as observed in the ScaNramp structure (Figures 4B and 7E) (Ehrnstorfer et al., 2014). In this conformation the Mn^2+^-coordination sphere would include four residues and two waters—analogous to the G223W structure—thus facilitating eventual metal release. The proposed transition from four to six to four Mn^2+^-coordinating residues could help preferentially stabilize the occluded transition state(Shilton, 2015) through the free energy (entropy-driven) gains of releasing the two water ligands. Furthermore, the rearrangements needed to achieve the hypothetical intermediate six-residue Mn^2+^ coordination—the helical extension and inward movement of TM6a, and the toppling of TM10’s top half—also close the external vestibule, providing a potential mechanistic link between local metal-coordination changes and bulk conformational change. To return to the outward-open state, the transporter must pass through an apo-occluded state as seen in the G45R structure, in which the N59 and Q378 sidechains reorient to stabilize D56 in the absence of the divalent cation carried during the outward-to-inward transition (Figures 4C and 7E). As the transporter reaches the outward-open state seen in EcoNramp (Ehrnstorfer et al., 2017) and our apo G223W structure, a protonation event at D56 may prime the binding site to receive another incoming metal ion. Future molecular dynamic simulations and/or experiments will be essential to test these predictions.

The mechanism described above for the DraNramp transport cycle—developed from structures of the same Nramp homolog in three distinct conformations and supported by metal and proton transport data—differs significantly from those previously observed for other LeuT-fold transporters. Mhp1, BetP, and (to a lesser degree) LeuT generally obey a “rocking bundle” model in which the rigid-body movement of four TMs that contain the primary substrate binding site (1, 2, 6, and 7) against the remaining TMs (3-5, 8-10) leads to conformational change (Forrest and Rudnick, 2009; Kazmier et al., 2017; Shi, 2013). In DraNramp TMs 4, 5, and 10 join TMs 1 and 6 to form the substrate-binding “mobile domain,” while TMs 2 and 7 join the remaining TMs as part of the scaffold. Furthermore, the mobile helices do not move as rigid bodies, as conserved helix-breaking motifs free TMs 1a, 6a, and the top of TM10 to move independent of TMs 1b, 6b and the bottom of TM10. In contrast, the fully-helical TM5 wholly reorients, and may thus coordinate the opening and closing of the inner and outer gates, connecting TMs 1a, 4, and 6b with TM6a (Figures 2 and 7A-B).

In comparison to other APC superfamily members, the large TM1a displacement in DraNramp most closely resembles its dramatic movement in LeuT(Krishnamurthy and Gouaux, 2012). Gating roles for TMs 5 and 10 have been ascribed for BetP, Mhp1, and MhsT (Malinauskaite et al., 2014; Ressl et al., 2009; Shimamura et al., 2010), although not as extensive as we propose here in DraNramp. Not surprisingly, the DraNramp conformational changes are most similar to those predicted by comparing structures of two other bacterial Nramp homologs in complementary conformations (Ehrnstorfer et al., 2014; Ehrnstorfer et al., 2017), suggesting conservation within the Nramp clade of the LeuT-fold family.

Whereas the distinct conformational changes of DraNramp demonstrate the diverse repertoire of dynamics available to the LeuT-fold family, the most striking mechanistic differences between DraNramp and other structurally-studied LeuT-fold transporters concern the coupling ion behavior. Most well-characterized members (including LeuT, BetP, and Mhp1) are Na^+^-driven symporters of small organic molecules which have one or two Na^+^-binding sites (Perez and Ziegler, 2013; Rudnick, 2013). Sodium binding at the highly-conserved Na2 site connects the “bundle” (TM1) and “scaffold” (TM8) domains while also shifting the conformational equilibrium to favor the outward-open state (Claxton et al., 2010; Tavoulari et al., 2016; Zhao et al., 2011). This Na2 site consists of hydroxyls from two consecutive serines/threonines on TM8 and four main-chain carbonyls (one from TM8, three from the unwound-region of TM1) (Perez and Ziegler, 2013; Yamashita et al., 2005). Intriguingly, the analogous location in DraNramp also contains highly-conserved hydroxyl-providing TM8 residues S327 (92%) and S328 (20%, 74% T), which may be remnants of the ancestral Na2 site conversion into a H^+^ site in the Nramp clade. This hypothetical evolutionary switch has precedent within the LeuT-fold family, as the proton-coupled amino acid transporter ApcT analogously uses a conserved TM5 lysine (K158), whose sidechain protrudes into the Na2 location, as its primary proton-binding site (Shaffer et al., 2009). The sodium-to-proton switch may have evolved in Nramps to avoid simultaneously coordinating two metal cations (Na^+^ coordination, like Mn^2+^ coordination, requires ∼6 oriented ligands, whereas H^+^ binding requires a single sidechain).

LeuT and other bacterial homologs also antiport a proton as they return to an outward-open state (Kantcheva et al., 2013; Zhao et al., 2010; Zomot et al., 2007) via a conserved glutamate (E290) on TM7 (Malinauskaite et al., 2016), analogous to the highly-conserved N275 (100%) that lines DraNramp’s intracellular vestibule. Available structures and MD simulations suggest that proton symport in ApcT and antiport in LeuT likely occur through the bulk opening and closing of the same permeation pathways used by the primary substrates (amino acids) (Krishnamurthy and Gouaux, 2012; Malinauskaite et al., 2016; Shaffer et al., 2009; Shi and Weinstein, 2010). In contrast DraNramp does H^+^ uniport even when mutationally (G223W) or chemically (A61C-MTSET) precluded from opening the intracellular vestibule (Figure 5). We propose a proton route from D56 through D131 and into a conserved salt-bridge network between TMs 3, 4, 8, and 9 (Figure 6), which remain relatively stationary during the conformational change process (Figure 2). Indeed, evolutionary analysis reveals that this polar network is unique to the Nramp clade of the LeuT-family (Cellier, 2016); this region is mainly hydrophobic in both LeuT and ApcT (Shaffer et al., 2009; Yamashita et al., 2005). A parallel transport pathway for protons alleviates the electrostatic problem of stabilizing three added positive charges (the proton and divalent metal cation) throughout the conformational change process. Such proton uniport requiring only subtle conformational rearrangements is more reminiscent of H^+^ shuttling in the CLC family of Cl^-^/H^+^ antiporters (Accardi and Miller, 2004; Accardi and Picollo, 2010; Basilio et al., 2014; Miller, 2006) than the canonical Na^+^ transport seen in LeuT-family symporters.

The DraNramp proton and metal transport mechanism, where primary and driving substrates enter via a common permeation pathway but exit via separate routes to the cytoplasm—with H^+^ transfer perhaps preceding and triggering bulk conformational rearrangement needed for Mn^2+^ release to occur—is thus far unique to the Nramp clade within the APC superfamily. This new model for Nramp transport therefore illustrates the evolutionary flexibility and adaptability of the shared LeuT fold.

## Methods

### Cloning of DraNramp

WT and mutant DraNramps were cloned in pET21-N8H as described (Bozzi et al., 2016b). All constructs were full-length, except the G223W crystallization construct was N-terminally truncated to residue 35; this deletion did not affect metal transport (Bozzi et al., 2016b). Mutations were made using the Quikchange mutagenesis protocol (Stratagene) and confirmed by DNA sequencing. Single-cysteine constructs also included the C382S mutation to remove the lone endogenous cysteine. The C41(DE3) *E. coli* strain was used for protein expression and *in vivo* assays.

### Expression and purification of DraNramp crystallography constructs

Six liters of DraNramp C41(DE3) cells were cultured as described (Bozzi et al., 2016b), pelleted and flash-frozen in liquid nitrogen. Proteins were purified at 4°C. Thawed cells were lysed by sonication in 40 ml load buffer (20 mM sodium phosphate, pH 7.5, 55 mM imidazole, 500 mM NaCl, 10% (v/v) glycerol) plus 1 mM PMSF, 1 mM benzamidine, and 0.3 mg/ml each DNAse I and lysozyme. Lysates were cleared for 20 min at 27,000 × *g*. Membranes were pelleted from supernatant at 230,000 × *g* for 70 min, homogenized in 65 ml load buffer and flash-frozen in liquid nitrogen. Thawed membranes were solubilized for 1 h, adding 1.5% (w/v) n-dodecyl-β-D-maltopyranoside (DDM), then spun at 140,000 × *g* for 35 min to pellet debris. Pre-equilibrated Ni-sepharose beads (3 mL; GE Healthcare) were incubated with the supernatant for 1 h, and washed with load buffer containing sequentially 0.03 % DDM, 0.5% lauryl maltose neopentyl glycol (LMNG), and 0.1% LMNG. Protein was eluted in 20 mM sodium phosphate, pH 7.5, 450 mM imidazole, 500 mM NaCl, 10% (v/v) glycerol, 0.01% LMNG, concentrated to < 0.5 mL in a 50-kDa cutoff centrifugal concentrator, and loaded onto a Superdex S200 10/300 (GE Healthcare) pre-equilibrated with SEC buffer (10 mM HEPES pH 7.5, 150 mM NaCl, 0.003% LMNG). Peak fractions were pooled, concentrated to ∼20-40 mg/ml, aliquoted, and flash-frozen in liquid nitrogen.

### DraNramp crystallization, X-ray diffraction data collection, and structure determination

Protein (10-15 μL) was loaded into a 100-μL glass syringe attached to an LCP coupling device (Formulatrix). A second 100-μL syringe containing 1.5 volumes of liquid monoolein (T > 37°C) was attached to the coupling device, and the two solutions were mixed for 100 cycles using an NT8 (Formulatrix) in LCP mixing mode at 5 mm/s. LCP boluses (50-100 nL) and precipitant (600-1000 μL) were dispensed into 96-well LCP glass plates and incubated at room temperature (RT). Crystals of G223W (25 mg/ml) with 2.5 mM MnCl2 in SEC buffer grown in 28% PEG400, 5 mM MnCl2, 100 mM MES pH 6, 50 mM succinic acid pH 6, 10 mM spermidine pH 7, (10-30 μm square plates) were harvested after nine days using mesh loops (MiTeGen) to scoop the bolus, then flash-frozen in liquid nitrogen. Similar G223W crystals in the apo state were obtained after seven days with 26% PEG400, 100 mM MES pH 6, 50 mM succinic acid pH 6, 20 mM spermidine pH 7. Crystals of the analogous NEM-modified G223C/C382S ΔN34 (G223C retains close to WT-level metal transport before NEM-labeling (Bozzi et al., 2016b)) were obtained in the same condition. Crystals of G45R (22 mg/ml) grown in 20% PEGMME 550, 150 mM NaCl, 100 mM HEPES, pH 7.0 (∼100 μm rods) were harvested after ∼10 days.

Data were collected at Advanced Photon Source beamline 24ID-C. Crystals were located by grid scanning with a 70-μm beam at 70% transmission followed by focused grid scanning with a 10-μm beam at 100% transmission. Data wedges were typically collected from -30° to +30° in 0.2° increments using a 10-μm beam at 100% transmission. Two wedges from 2 crystals (G45R), 23 wedges from ∼15 crystals (G223W with Mn^2+^), or 39 wedges from ∼20 crystals (G223W apo) were independently indexed and integrated then combined during scaling using HKL2000 (Otwinowski and Minor, 1997) to obtain complete datasets. Structures were determined using software provided by SBgrid (Morin et al., 2013). Initial phases were obtained by molecular replacement in PHASER (McCoy et al., 2007) using our first DraNramp structure (PDB: 5KTE) as a search model for G45R and an in-progress G45R model for G223W. Model building and refinement were iterated in Coot(Emsley and Cowtan, 2004) and PHENIX (Adams et al., 2010), respectively. For all structures, positional and B-factor refinement with TLS restraints were used throughout, with torsion angle and NCS restraints for G45R, and secondary structure restraints for the Patch mutant. G45R contains two protein molecules in the asymmetric unit—chain A with residues 45-167 and 174-436 and chain B with residues 44-168 and 175-436 (RMSD 0.86 Å over 2899 atoms, 0.50 Å over all 386 Cαs)—and seven fully or partly modeled monoolein molecules. Chain A was used for figures and analyses. The G223W Mn^2+^-bound structure includes residues 39-436, nine full or partial monooleins, one spermidine molecule, and two Mn^2+^ ions—one in the metal-binding site, one at a crystal-packing interface. The G223W apo structure includes residues 39-436 and six full or partial monooleins. The two G223W structures align with RMSD 1.41 Å over 3012 atoms, 1.08 Å over all 398 Cαs). The electron density for each TM is shown in Figure Figure 2—figure supplement 2A for G45R and Figure 2—figure supplement 2B for G223W with Mn^2+^, while Figure 2—figure supplement 2C shows the metal-binding site for both the Mn^2+^-bound and apo G223W structures. The inward-open Patch mutant structure was updated to correct the position of intracellular loop 10-11 and the registry of TM11, and extend the N-termini of TMs 5, 7 and 9, the C-terminus of TM7, and extracellular loop 7-8, and improve the geometry of the Fab. The new model comprises residues 43-165, 170-236, 256-341, 351-436 of DraNramp, 1-129 and 132-213, and 1-213 of the Fab heavy and light chains, respectively, and three Os^3+^ ions.

### In vivo metal transport assays

Metal uptake assays in *E. coli* were performed as described previously (Bozzi et al., 2016a). For each biological replicate reported in figure legends, a separate culture of transformed *E. coli* was grown and induced to express the exogenous Nramp construct.

### In vitro transport assays

Protein purification, liposome preparation, and metal/proton transport assays were performed as described in an accompanying paper (Bozzi et al., 2018). Single-cysteine constructs A53C and A61C were purified in the presence of 1 mM DTT. To modify cysteines, liposomes were diluted into buffer (120 mM NaCl, 10 mM MOPS pH 7) containing 3 mM MTSET, 3 mM MTSEA, or 4 mM NEM, and incubated at least 30 min at RT before beginning transport assays. For each technical replicate reported in figure legends, a separate aliquot of dye-loaded proteoliposomes was diluted into the appropriate outside buffer, including cysteine modifiers if applicable, then fluorescence time course data were collected before and after addition of valinomycin, metal substrate, and/or ionomycin.

### Cysteine accessibility measurements

Cells grown as for the uptake assay were washed once in labeling buffer, resuspended at OD600=2, and aliquoted 100 μL per well in a 96-well plate. A 1:1 NEM dilution series was prepared in labeling buffer at 8 mM; 100 μL of the appropriate 2X NEM solution was added to each well and incubated 15 min at RT. L-cysteine (10 μL of 200 mM) was added to quench NEM. Cells were washed twice in labeling buffer, pelleted, resuspended in 30 μL lysis and denaturing buffer (6 M urea, 0.1% SDS, 100 mM Tris pH 7) with 0.5 mM DTT and incubated 1 h at 37°C. The lysate (10 μL) was mixed with 3.5 μL of 6 mM 5K-PEG maleimide (Creative PEGWORKS) in lysis and denaturing buffer, incubated 1 h at 37°C, and the reaction terminated by adding sample buffer with β-mercaptoethanol. Protein was detected via SDS-PAGE and Western blotting using an Alexa 647-conjugated anti-His-tag antibody (QIAGEN) and a Typhoon Imager (GE Healthcare). ImageJ64 was used to determine the % modification as described (Bozzi et al., 2016b). For each biological replicate reported in figure legends, a separate culture of transformed *E. coli* was grown and induced to express the exogenous Nramp construct.

### Sequence alignments

An alignment of 9289 Nramp sequences was obtained from a HMMER (Finn et al., 2011) search using the DraNramp sequence with an E-value of 1, filtered for sequences with just one domain, then filtered for sequences 400-600 residue long. Incomplete sequences and sequences lacking the canonical Nramp TM1 “DPGN” and TM6 “MPH” motifs were removed, yielding 6878 sequences. A seed of 92 diverse sequences were aligned using MUSCLE (Edgar, 2004), then HMMER was used with a gap threshold of 0.99 to create the final alignment (Figure 2—source data 1).

### Structural comparisons and analyses

Per-residue Cα RMSD values were calculated using the ColorByRMSD PyMOL script, and whole-structure Cα RMSD values using the PyMOL align command with cycles=0. To generate the distance difference matrices, the pairwise distances between all Cα atoms were calculated for each structure (inward, inward-occluded, and outward). Then, for each combination of two conformations, a distance difference matrix was calculated by taking the difference between the distance matrices corresponding to each conformation. These distance-difference values were then averaged for each pair of TM helices (after dividing TMs 1, 6, and 10 into “a” and “b” segments) using an RMSD-like calculation to obtain a 14×14 matrix for each pair of conformations. pK_a_ values were calculated using PROPKA (Dolinsky et al., 2004) with CHARMM forcefields.

## Data availability

The accession number for the DraNramp crystal structures reported in this paper are G45R inward-occluded, PDB ID: 6C3I; G223W ΔN34 outward-open with Mn^2+^, PDB ID: 6BU5; G223W ΔN34 outward-open apo, PDB ID: 6D91; revised inward open, PDB ID: 6D9W. The unprocessed diffraction images were deposited in the Structural Biology Data Grid (https://data.sbgrid.org/) with SBDG ID: 567 (G45R); 564 and 576 (G223W ΔN34 Mn^2+^-bound and apo respectively); and 332, 333, and 334 (inward-open). The raw biochemical data that support the findings of this study are available from the corresponding author upon reasonable request.

## Author Contributions

R.G. oversaw and designed the research with A.T.B. and C.M.Z.; A.T.B. performed all functional assays and analyzed the resulting data; B.K.L. designed, cloned, and initially validated the panel of tryptophan mutants with A.T.B.; C.M.Z, obtained crystals and diffraction data and solved the structure of G45R DraNramp; A.T.B. obtained crystals and diffraction data and solved the structures of G223W ΔN34 DraNramp, with assistance at each stage from J.M.N., C.M.Z., and/or R.G.; R.G. corrected the registry and re-refined the original DraNramp structure; C.H.Z. computationally analyzed Nramp sequences and structures. A.T.B., C.M.Z., and R.G. wrote the manuscript, with input from all authors.

## Acknowledgements

We would like to thank Andrew Kruse for assistance with LCP crystallography, Anthony Hesser for initial DraNramp crystallization, and Chris Miller, Joe Mindell, Niels Bradshaw and Gaudet lab members for helpful discussions. This work was funded in part by NIH grant 1R01GM120996 (to R.G.) and a Jane Coffin Childs Postdoctoral Fellowship (to C.M.Z.). We thank the NE-CAT beamline staff at the Advanced Photon Source (Argonne, IL, USA) for help with data collection. NE-CAT is funded by NIH (P41 GM103403 and S10 RR029205), and the Advanced Photon Source by the U.S. Department of Energy (DE-AC02-06CH11357).

---

**Figure 1—figure supplement 1.**
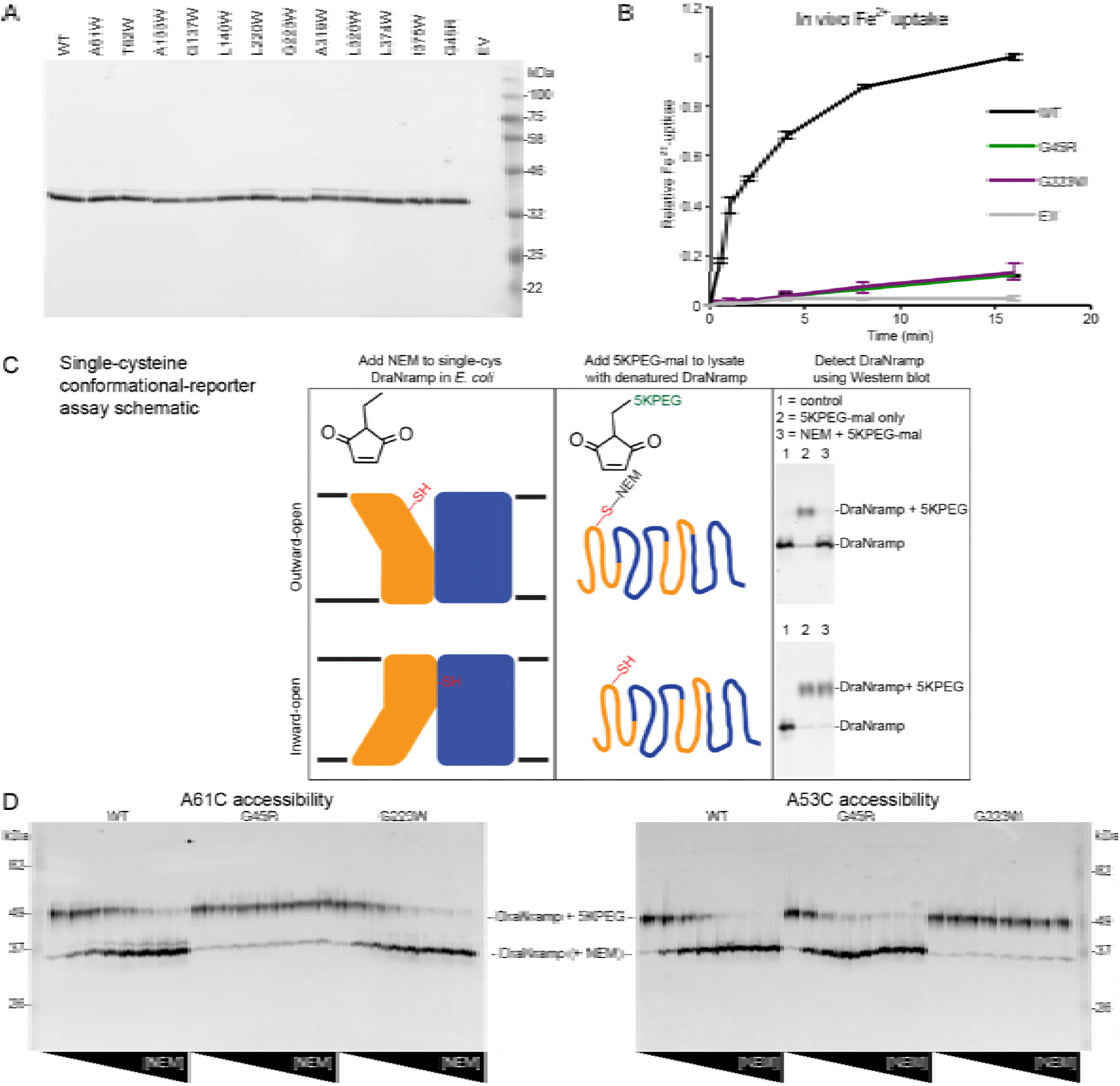
Validation of conformationally-locked crystallization constructs. (A) Western blot against N-terminal His-tag for total lysates show most tryptophan mutants expressed similarly to WT DraNramp in *E. coli*. (B) Both G45R and G223W eliminated Fe^2+^ uptake in *E. coli*. (C) Schematic for cysteine accessibility assay two-step protocol using conformational reporters: first a gradient of NEM was applied to label any solvent-accessible cysteines, then the cells were lysed and protein denatured and 5K-PEG maleimide was added, which reacted with any previously unmodified cysteines. The 5K-PEG maleimide-labeled protein shifted upward from the NEM-labeled protein on SDS-PAGE. (D) G45R and G223W had opposing effects on single-cysteine reporter accessibility to NEM modification in *E. coli*. The A61C reporter—only solvent-exposed in the outward-facing state—was fully protected from NEM-labeling by the G45R mutation, indicating an inward-locked state. In contrast, A53C— likely only accessible in an inward-facing conformation—was fully protected from NEM-labeling by G223W, indicating an outward-locked state. Both reporters were fully labeled at high NEM concentrations for WT, suggesting the entire protein population cycles through both conformational states during the NEM labeling time.

**Table 1—figure supplement 1.**
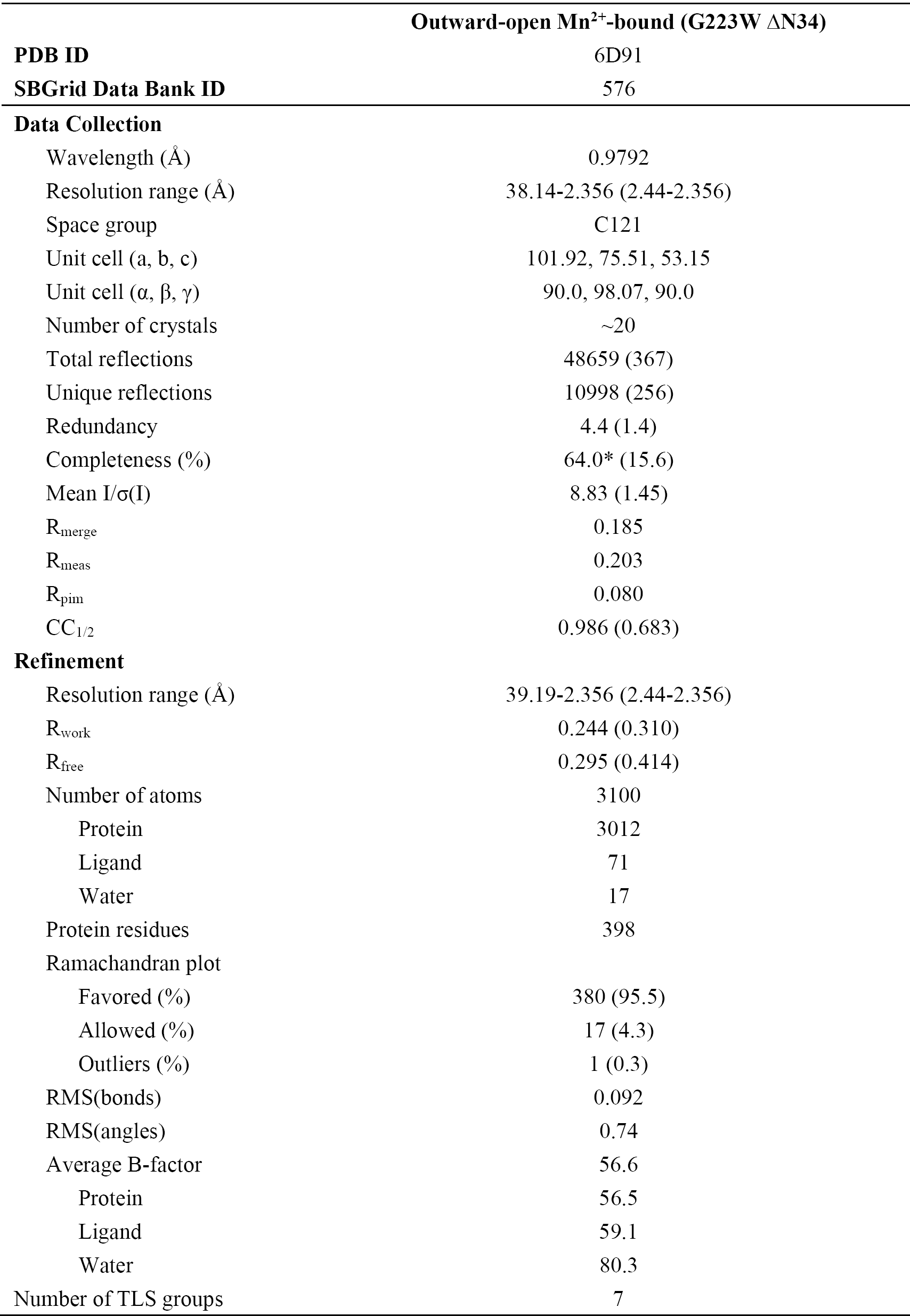
Data collection and refinement statistics for apo G223W ΔN34.

**Figure 2—figure supplement 1.**
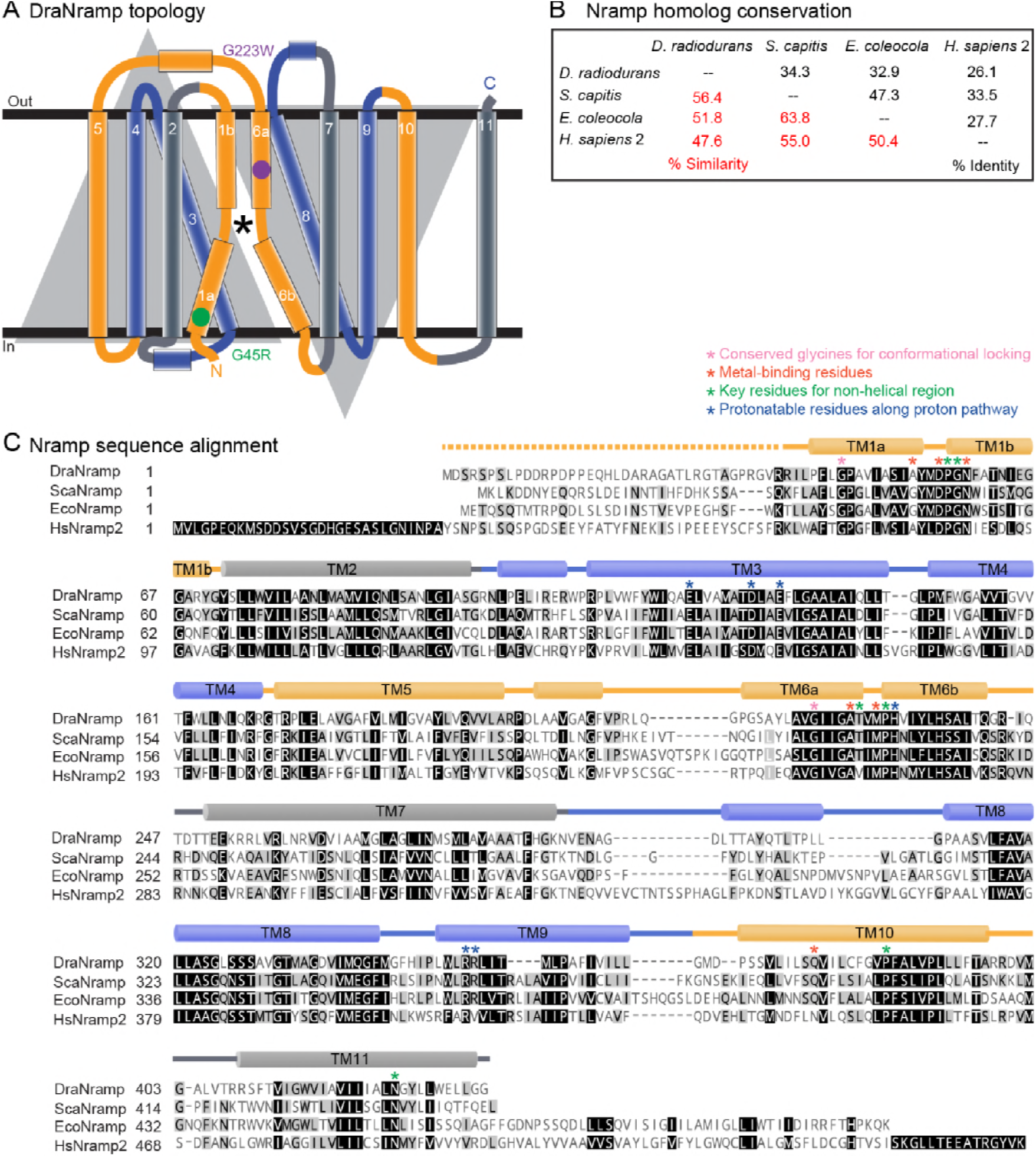
Secondary structure and primary sequence of DraNramp. (A) DraNramp consists of 11 TMs, with the first ten TMs arranged in the canonical LeuT-fold: two pseudosymmetric inverted repeats of five TMs intertwine in the tertiary structure. The two conformational-locking mutations, G45R and G223W, on TMs 1a and 6a respectively, are at analogous positions within each repeat. The first and last TM in each repeat (1 and 5, and 6 and 10; gold) move significantly between the G223W and G45R structures, while the middle three remain relatively stationary (2-4, and 7-9; gray or blue). TMs 1, 4, 5, 6, and 10 group into a mobile domain, with the remaining six TMs considered a scaffold domain. ^*^ denotes the metal-binding site. (B) Sequence identity and similarity of DraNramp with *S. capitis* (Sca) and *E. coleocola* (Eco) Nramp homologs, which have been crystallized in the inward-facing^24^ and outward-facing states^31^ respectively, and human Nramp2 (DMT1). Sequences were aligned using the BLOSUM62 matrix, with a cutoff of 1 used for the reported % similarity. (C) Sequence alignment of DraNramp, ScaNramp, EcoNramp, and HsNramp2 shows high conservation at positions of apparent structural significance and demonstrated functional importance.

**Figure 2—figure supplement 2.**
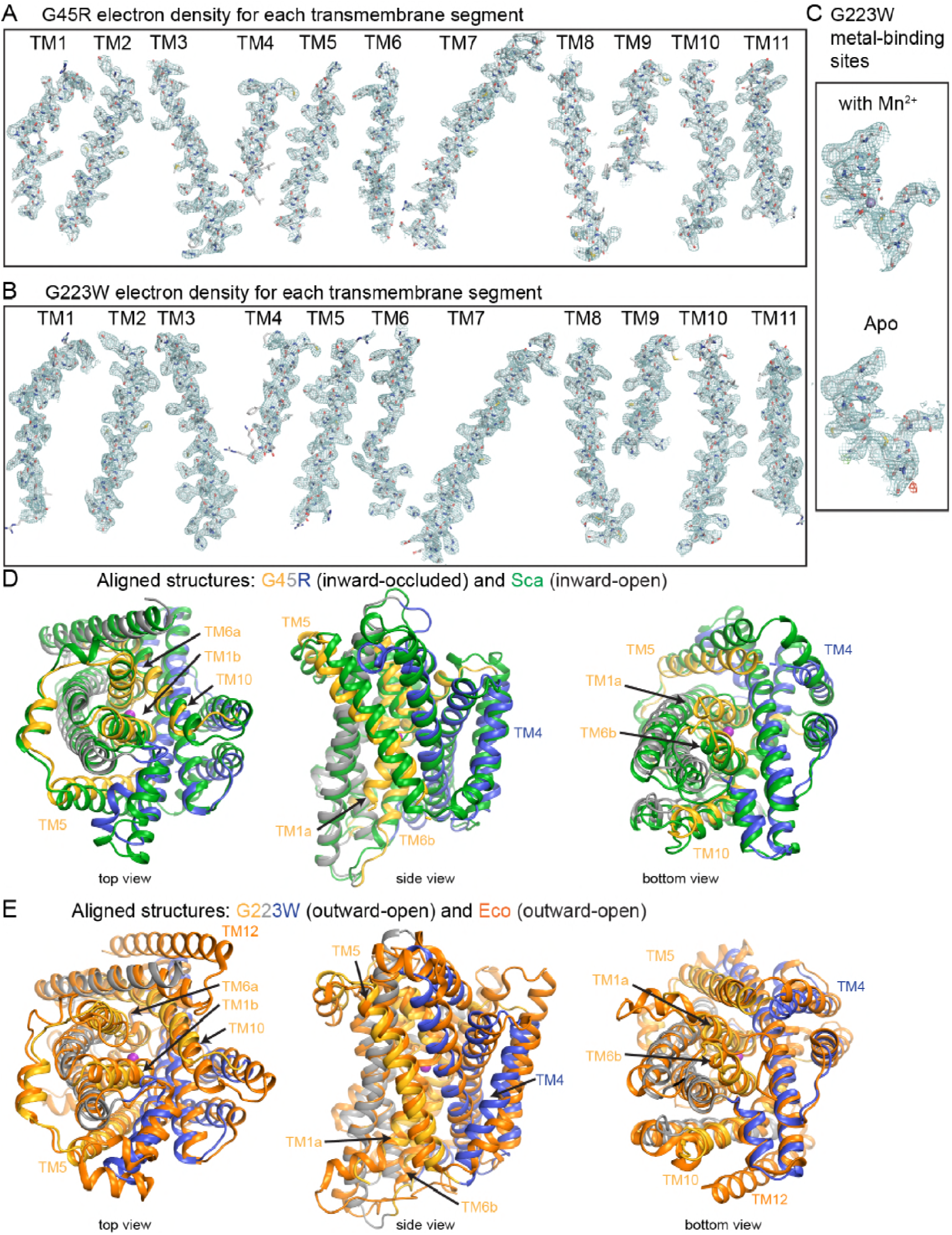
Electron density maps for new crystal structures and structural comparisons to other Nramp homologs. 2F_o_-F_c_ electron density maps for each transmembrane segment of (A) G45R (6C3I) and (B) G223W (6BU5) structures contoured at 1.0 σ. (C) Electron density maps for the metal-binding site for the Mn^2+^-bound (6BU5) and apo (6D91) structures, with the latter also showing the F_o_-F_c_ difference maps contoured at +/- 3.0 σ (green/red). Superpositions of (D) G45R with ScaNramp (green) and (E) G223W with EcoNramp (orange) confirm that our new structures represent outward-open and inward-occluded conformations, respectively. The views in (D) and (E) are based on a superposition of the G45R and G223W structures.

**Figure 2-source data 1. Nramp sequence alignment.**

Alignment of 6878 Nramp sequences generated with HMMER and filtered to contain only sequences with canonical Nramp TM1 “DPGN” and TM6 “MPH” motifs.

**Figure 4—figure supplement 1.**
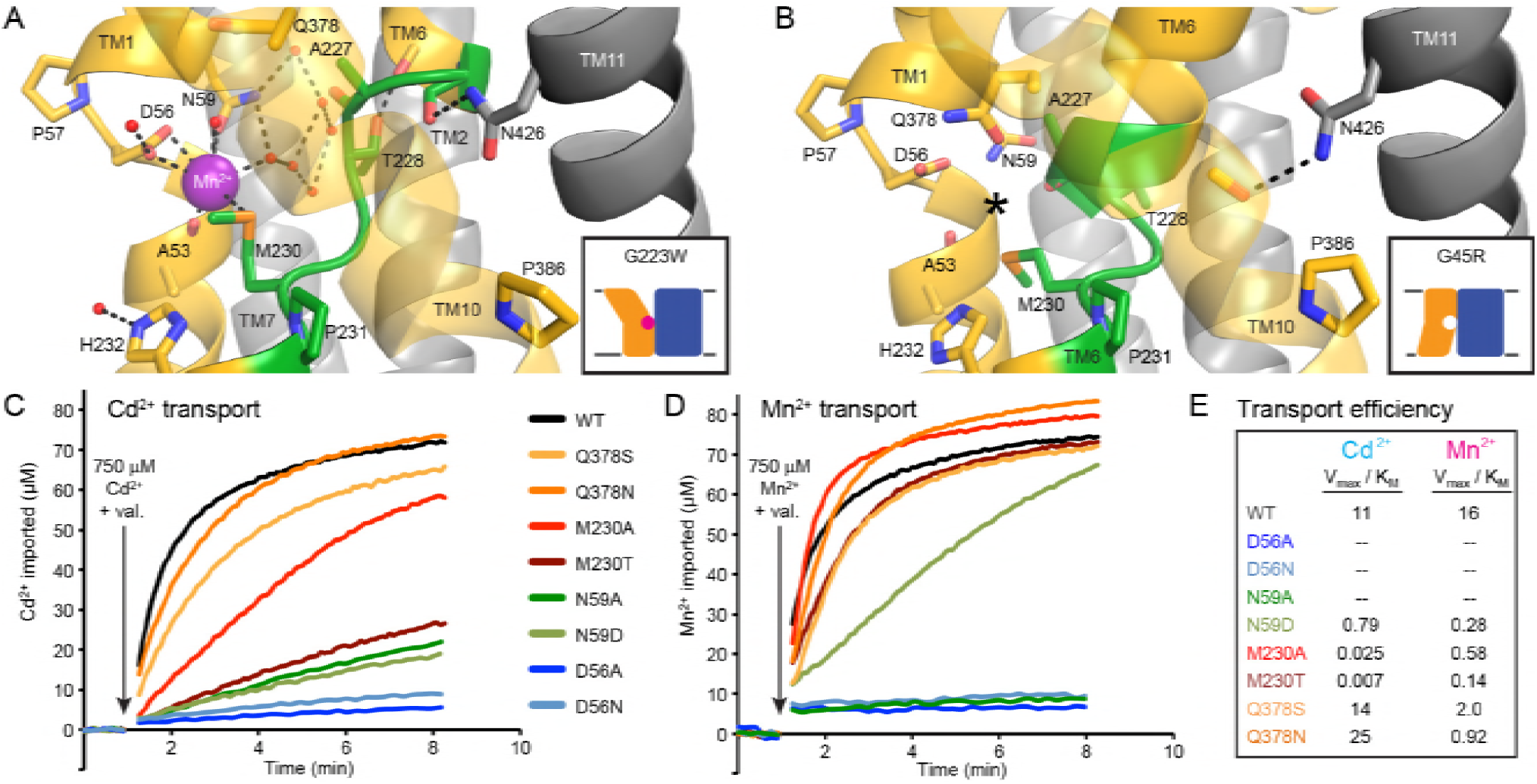
Extended TM6 unwound region enables metal-binding in outward-facing DraNramp. (A) TM6 unwinds in the outward-facing G223W structure (non-helical region encompassing residues 226-231 is shown in green). Conserved sidechains on TM6 (T228) and TM11 (N426) donate hydrogen bonds to backbone carbonyls on TM6a to stabilize the extended non-helical region. A network of ordered water molecules forms the outer coordination sphere for the bound Mn^2+^ and extends into the outward metal-permeation pathway between TMs 1b, 6a, and 10. Conserved H232 below the metal-binding site also coordinates an ordered water. (B) For comparison the same region is highlighted in green in the G45R inward-occluded structure, although the TM6a helix is extended, and only residues 228-231 are unwound. Interestingly, in this structure the N426 sidechain donates a hydrogen bond to satisfy the backbone carbonyl on TM10 just above the conserved P386. For clarity, TMs 3, 4, 5, 8, and 9 are omitted and TM10 is shown as a transparent ribbon in panels A and B. (C-D) Representative traces (n = 3) for WT and binding-site mutants (C) Cd^2+^ and (D) Mn^2+^ uptake at 750 μM. Mutations to D56 eliminate all metal transport, while mutations to N59 severely impair transport for both metals. At higher metal concentrations, mutations to M230 are much more deleterious to Cd^2+^ transport than Mn^2+^ transport, while mutations to Q378 do not greatly impair transport of either metal. (E) Transport efficiency (V_max_/K_M_) comparisons from the data in Figure 3E-F illustrate all tested mutations are deleterious to Mn^2+^ transport, while all but Q378S and Q378N are deleterious to Cd^2+^ uptake.

**Figure 5—figure supplement 1.**
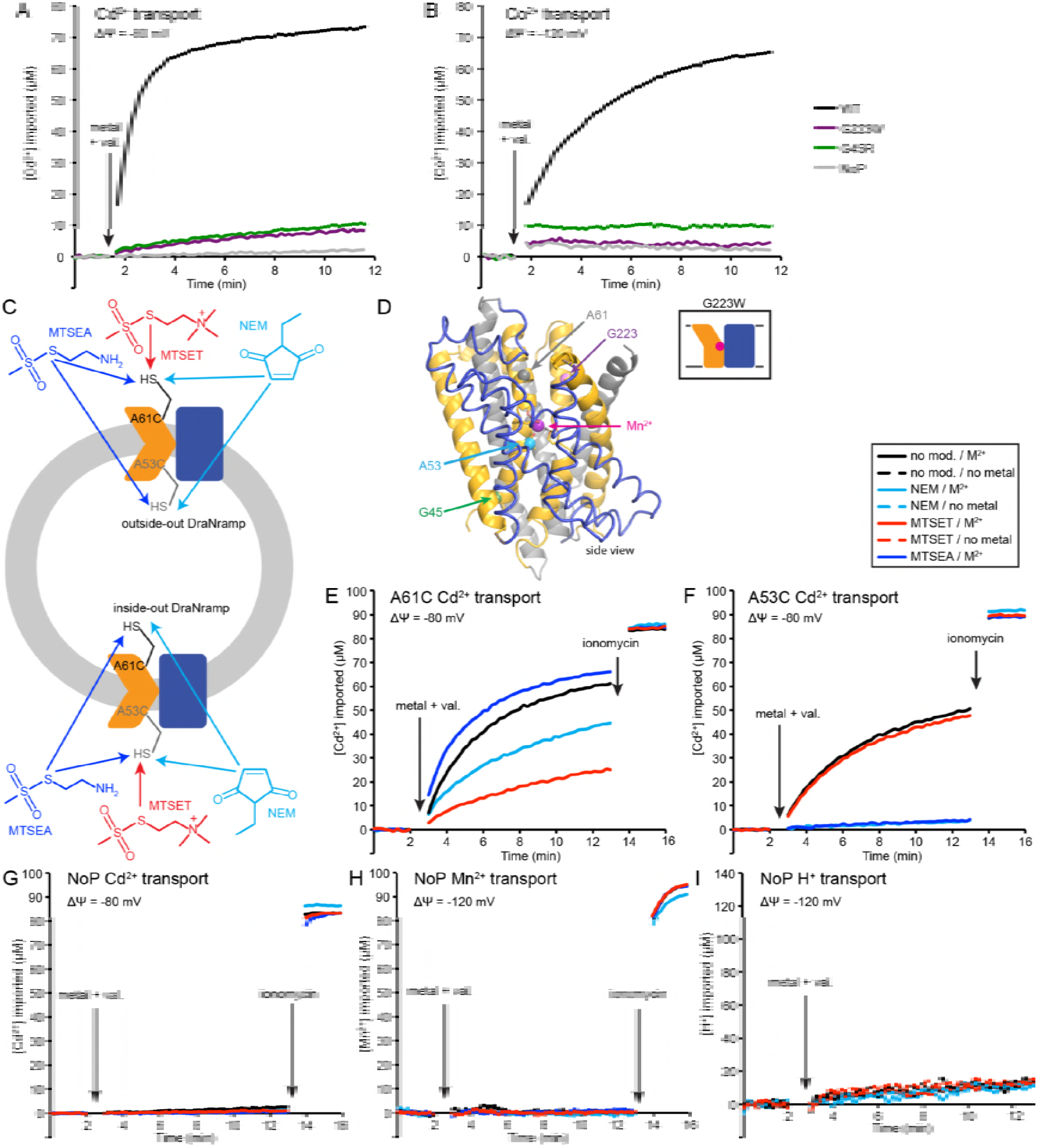
Conformational locking impairs metal transport. Sample traces (n=4) of (A) Cd^2+^ and (B) Co^2+^ transport over time in proteoliposome. WT enabled robust uptake while G45R and G223W did not. NoP = No Protein control liposomes. (C) Schematic for *in vitro* cysteine modification of A53C (inside accessible) and A61C (outside accessible). Assuming random insertion into liposomes, there is a ∼50:50 mix of right-side-out and inside-out transporters. A53C and A61C should thus be fully-labeled by membrane-permeable MTSEA and NEM for transporters in both orientations. In contrast, only right-side-out A61C and inside-out A53C transporters should be labeled with the membrane-impermeable MTSET. (D) A61 and G223 line the metal permeation pathway to binding site, while A53 and G45 face the metal exit pathway to the cytoplasm. (E,F) Sample traces (n = 3) of Cd^2+^ transport over time in the presence or absence of cysteine modifying agents, with trends similar to Mn^2+^ transport in Figure 5E,G: (E) MTSET greatly reduced A61C Cd^2+^ transport, NEM had a moderate effect, and MTSEA had a slight enhancing effect; (F) NEM and MTSEA eliminated A53C Cd^2+^ transport, while MTSET had no effect. (G-I) Sample traces (n = 3) show that none of the modifiers enabled M^2+^ or H^+^ leak into empty liposomes.

**Figure 6—figure supplement 1.**
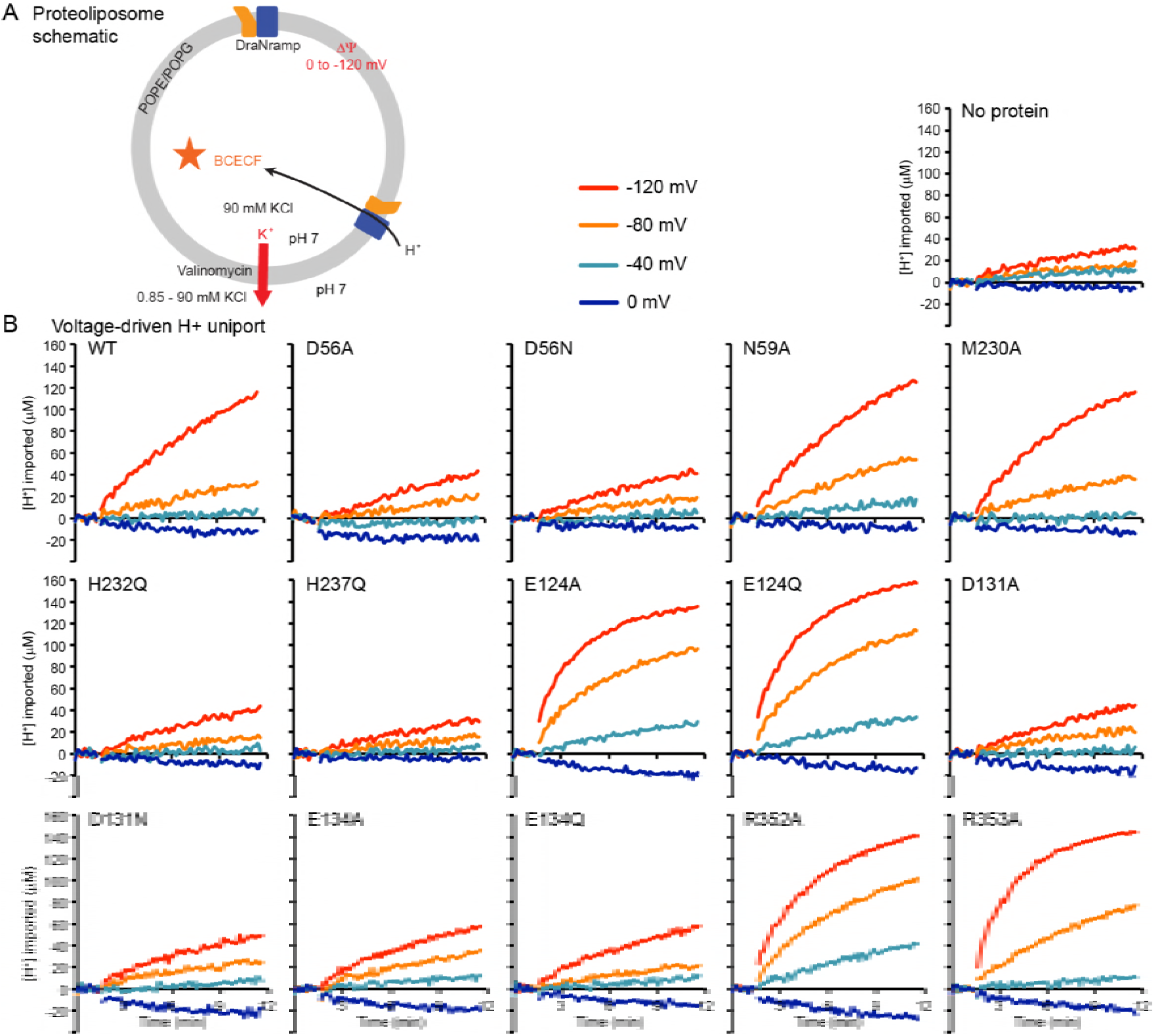
Sample traces illustrate mutant perturbations of voltage-driven proton uniport. (A) Schematic for proteoliposome proton uptake assay. (B) Representative time traces (n ≥ 4) of H^+^ uptake into liposomes measured at four Δψ values for each mutant or no-protein control.

**Video 1. Internal rearrangements during DraNramp conformational changes.**

Morph of the structure of DraNramp based on a global superimposition of the three captured conformational states. TMs 1, 5, 6, and 10 are colored gold, TMs 3, 4, 8, and 9 blue, and TMs 2, 7, and 11 gray, viewed in the membrane plane and the intracellular face pointing down. The morph starts from the G223W outward-facing conformation, transitions to the G45R occluded state, then to the patch mutant inward-open state. The view then shifts to the intracellular face, followed by the external face, continuing to alternate back and forth through these three conformations.

